# Cell organization and disturbance-mapping using image activated cell-profiling (CODIAC)

**DOI:** 10.64898/2026.08.06.742695

**Authors:** Melanie E. S. Krause, Dan Lou, Shengdi Li, Haeyeon Jung, Daniel Kirrmaier, Krisztina Gubicza, Daniel Schraivogel, Michael Knop, Lars M. Steinmetz

## Abstract

Protein localization and abundance are tightly regulated to maintain cellular homeostasis, and their dysregulation is a hallmark of disease. However, methods that monitor the spatial localization changes of many proteins at once, and at scale, remain limited. Here we introduce CODIAC (Cell Organization and Disturbance-mapping using Image-Activated Cell-profiling), which couples image-enabled cell sorting (ICS) with improved Cas12a-assisted endogenous PCR tagging and machine-learning-guided gating to profile pools of fluorescently tagged proteins. Rather than isolating discrete cell populations, CODIAC reads the “phenotypic fingerprint” of each fluorescently tagged protein, defined by its distribution across a fixed set of image-defined sort bins. The method resolves localization-associated phenotypes, reproducibly profiles complex pools, generalizes to proteins absent from the training set, and detects shifts in protein localization and abundance following chemical perturbation. CODIAC thus extends ICS from outlier screening toward systematic profiling of the spatial proteome and its remodeling under perturbation.

## Introduction

Protein function depends on subcellular localization, which is dynamically regulated and frequently disrupted in diseases^1^. Fusion proteins expressed from endogenous loci are widely used to study protein localization because they preserve native expression levels and avoid overexpression artifacts. Such fluorescently tagged proteins are conventionally analyzed by fluorescence microscopy, which offers high-resolution spatial information^2^ but cannot isolate cells with specific phenotypes at scale. Advances in gene editing tools have improved the precision and scalability of endogenous tagging in mammalian cells^3,4^, and enabled large-scale parallel tagging through optimized split-protein approaches^2,5^. As tagging itself becomes scalable, the limiting step shifts downstream to how many tagged proteins can be read out at once. Two factors constrain this: the spectral bandwidth of fluorescent reporters restricts simultaneous observation to a few proteins per cell^6^, a limit that multiplexed immunofluorescence circumvents only in fixed cells^7^, and the per-line cost of generating and culturing tagged mammalian lines imposes a throughput bottleneck. Proteomics offers a complementary route to protein interactions and compartment composition, but yields limited information at single-cell resolution in high-content formats^8–10^.

Pooling differently tagged cells removes the need to culture and image each cell line separately, but in turn requires each imaged phenotype to be linked back to the corresponding tagged gene. Typically performed via *in situ* sequencing of barcodes, this methodology relies on complex microfluidic sample handling and repeated re-imaging^11–14^. An alternative strategy forgoes barcode sequencing altogether, using each tagged protein’s own localization pattern and expression level as a visual barcode to identify clones within multicolour pools by high-throughput microscopy and machine learning, and to track perturbation-induced localization changes^3,15^. Such image-based readouts are powerful for live-cell monitoring but remain bounded by microscopy throughput and resolve protein identity optically rather than through a sequencing-based measurement. Related multimodal efforts have built on resources such as the OpenCell Atlas^10^. An alternative to *in situ* genotyping is to sort cells by phenotype and identify the tagged gene afterward, by sequencing reporter insertion sites that act as endogenous barcodes^16,17^.

Image-enabled cell sorting (ICS) enables such a first-sort-then-sequence workflow. Unlike conventional flow cytometry, ICS captures image-associated parameters such as signal size, shape or correlation with a secondary marker, providing information beyond cell size, shape and signal intensity. ICS is part of a broader class of image-based cytometry and sorting technologies that capture either subcellular fluorescence patterns^18–30^ or non-fluorescent morphological features^31,32^. To date, ICS has served as a screening tool that enriches rare cells from a large population, typically based on the signal of a single labeled protein^26,33–36^. Yet the same spatial measurements of ICS should, in principle, enable quantitative profiling of complex pools of endogenously tagged proteins and to track how their localization shifts under perturbation, a capability that has not been demonstrated.

Here we establish CODIAC (Cell Organization and Disturbance-Mapping using Image-Activated Cell-Profiling), which couples ICS with pools of Cas12a-assisted endogenous tagged mammalian proteins. Applied to a reference pool generated from individual tagging of proteins that localize to major subcellular compartments, CODIAC resolves protein localization at scale, detects perturbation-induced phenotypic shifts, and establishes a machine-learning-guided strategy for sorting complex fluorescent cell populations. In doing so, it extends ICS from outlier screening to systematic profiling of the mammalian spatial proteome and of how it is remodeled under perturbation.

## Results

### Design of CODIAC

CODIAC infers the fluorescent phenotype of a tagged protein from its measurable ICS sorting outcome. Conventional sorting optimizes one set of gating parameters and thresholds, hereafter gating rules, to enrich a single target population. CODIAC instead assays pooled, fluorescently tagged cells in a multiplexed manner by evaluating their behaviour across a single, unified set of gating rules, translating morphological and fluorescent variation between cells into quantitative, sequencing-based readouts (Fig. 1a). The rule set is built to be universal across proteins. We therefore define a reference panel of proteins spanning diverse subcellular localizations, the training tag pool, and use it to establish the rules. Sorting this pool by ICS yields a series of bins of distinct composition, and maps each protein’s ICS outcome, its distribution across bins, to a fluorescent phenotype (Fig. 1b). Shifts in this distribution between conditions report phenotypic change upon perturbation (Fig. 1c).

**Figure 1:**
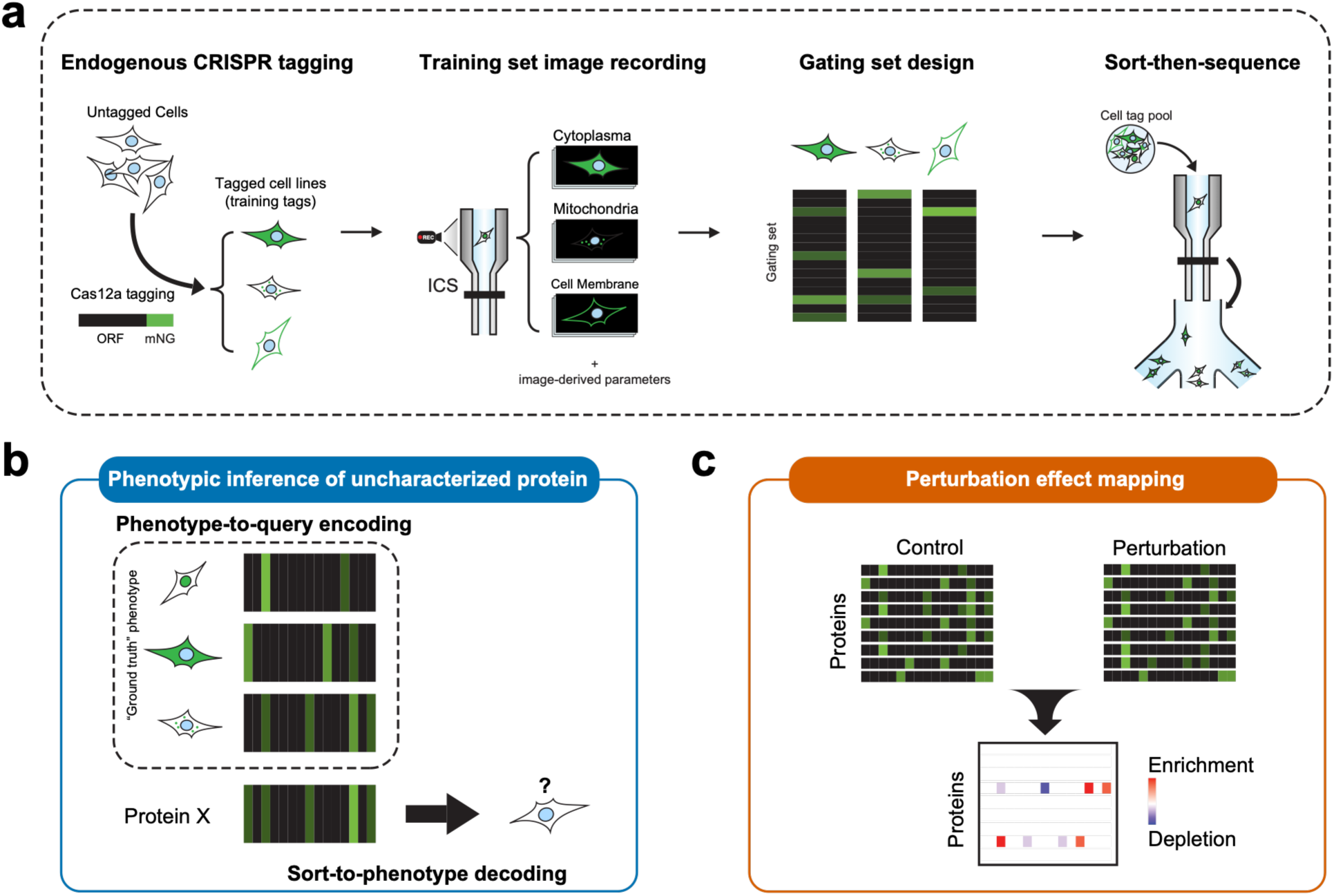
The CODIAC workflow. (1) Hek293T-N cells are endogenously tagged with mNeonGreen (mNG) at individual gene loci. (2) Individual populations are recorded on the BD S8 image sorter generating the ground truth training dataset. (3) Machine-learning powered query discovery then identifies optimal gating strategies, which (4) are applied to a complex pool. Sort bins are then sequenced to link genotype to sort bin phenotype. (b) A protein’s distribution across bins reflects its fluorescent phenotype, enabling phenotypic inference of uncharacterized proteins. (c) Comparison of sorting distribution shifts between perturbed and unperturbed conditions allows quantitative assessment of perturbation effects on fluorescent phenotypes.

### CRISPR-Cas12a-assisted PCR tagging enables efficient fluorescent cell line generation

To implement CODIAC, we established a scalable endogenous tagging strategy for building libraries of fluorescently tagged mammalian cells. We adapted PCR-based endogenous tagging^37,38^ in which a PCR-generated cassette carries homology arms supplied by the PCR oligos, the tag (mNG)^39^, and a 2A-linked selection marker^40^.

After antibiotic selection, flow cytometry showed a progressive increase in mNG-positive cells, reaching 30–54% for most genes (Fig. 2a, Supplementary Fig. 1a & 1b). To test performance on difficult targets, we generated cassettes for genes with only 2–15% mNG-integrated alleles in the OpenCell library^2^. After 15–18 days of selection, over 60% of cells showed mNG fluorescence at the expected subcellular location for 13 of 16 genes (Fig. 2b). Thus, our approach efficiently generates and enriches correctly localized mNG-tagged cell populations, including for genes previously considered difficult to tag.

**Figure 2:**
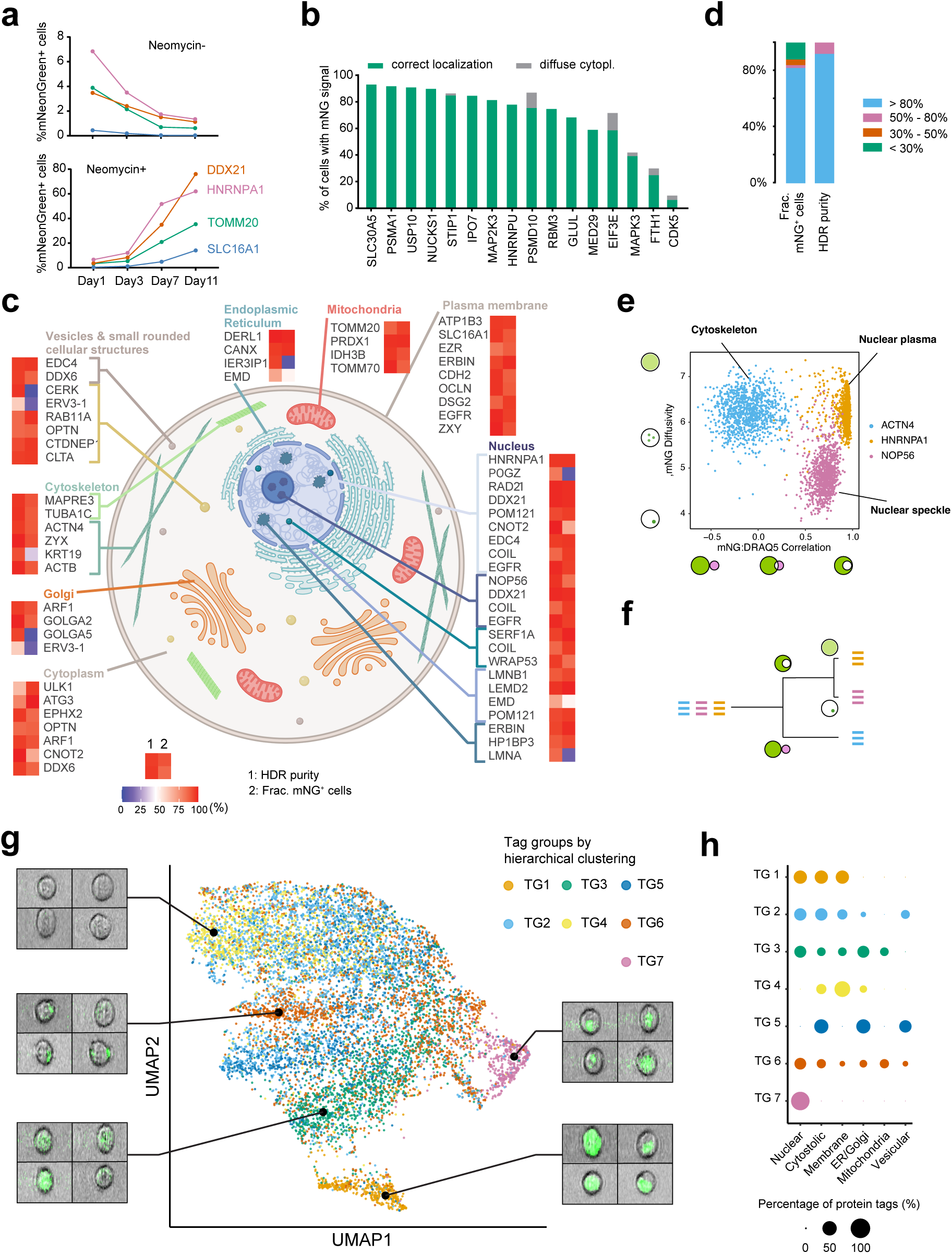
Cas12a PCR tagging enables the creation of large fluorescent protein libraries. (a) mNG⁺ cell fractions measured by flow cytometry over 11 days after tagging at several loci, (top) without and (bottom) with antibiotics selection. (b) Cells transfected with PCR cassettes targeting genes previously reported to have low tagging efficiency. Fluorescence microscopy of HOECHST-stained cells was used to determine the fraction of cells (in %) with mNG fluorescence at correct localizations. (c) Schematic illustration of a 50-protein library with targets spanning diverse subcellular localizations and quantification of cells with mNG signal at expected localizations via fluorescence microscopy. The efficiency of HDR-mediated editing (HDR purity) was measured using Anchor-Seq. (d) Characterization of editing efficiency for tagged genes in (c). The proportion of correctly localized mNG+ cells was quantified via fluorescence microscopy. HDR-mediated editing efficiency (HDR purity) was determined by Anchor-seq. (e) FACS plot displaying 3 protein tags plotted based on mNG signal diffusivity and correlation with DRAQ5. (f) Simplified decision tree modeling to separate three distinct protein phenotypes in (e). (g) UMAP visualization of 10,000 cells from 7 protein tag groups via mNG-relevant image parameters. Tag groups are derived from hierarchical clustering of 50 protein tags, setting cluster number equals to 7. Cell pictures are original image output from ICS, overlaying the brightfield (grey) and mNG (green) channels. (h) Distribution of protein localization for each tag group. Percentages represent the fraction of a tag group with specific localizations. Tags with multiple localizations are counted once per localization.

### Construction and characterization of a reference protein library

To understand how ICS can interpret complex subcellular phenotypes across a broad range of protein species, we tagged individually 50 genes in HEK293T cells representing the major subcellular localizations (Fig. 2c). Genes were selected based on high expression across cell types using the Human Protein Atlas and the OpenCell database^2^, with at least three genes representing a compartment. By fluorescence microscopy, 41 of 50 tags (82%) showed mNG at the correct localization in more than 80% of cells (Fig. 2d), and homology directed repair (HDR)-mediated editing efficiency (quantified by Anchor-Seq^16,37^ with a custom pipeline^38^) exceeded 50% for 46 tags (92%), with only one below 30% (Fig. 2d). Cell lines were then independently profiled by ICS to acquire single-cell fluorescence images (Supplementary Fig. 2a).

Beyond raw signal intensity, ICS extracts image-derived metrics describing fluorescence-signal morphology, which form the key feature space for our sorting-program design. Alongside conventional intensity measurements (area, height, width), the system records spatial descriptors of signal distribution (signal size, eccentricity, radial moment, diffusivity, delta center of mass, and correlation with a second imaging parameter). These parameters distinguish signals that are inseparable by conventional flow cytometry. For example; the nuclear proteins NOP56 and HNRNPA1 (Supplementary Fig. 2b) are separated from the cytoskeletal protein ACTN4 by the correlation between mNG and nuclear DRAQ5 signal (Fig. 2e). Among nuclear proteins NOP56 is further distinguished from HNRNPA1 by mNG signal diffusivity, reflecting distinct nuclear-speckle versus plasma localization patterns. Combining these parameters thus allows specific proteins to be enriched from a mixed pool (Fig. 2f).

Using the ICS data, we built a UMAP of cells across all 50 lines. This revealed a structured organization of proteins in ICS feature space according to their fluorescence phenotypes (Fig. 2g). Hierarchical clustering defined seven tag groups, each enriched for particular combinations of localizations (Fig. 2h; Supplementary Fig. 3). Notably, individual localization categories were distributed across multiple groups rather than mapping one-to-one onto a single group. This aligns with the expectation that proteins sharing a compartment can display distinct phenotypes shaped by expression level and co-localization across compartments, which together shape their imaging signatures (Supplementary Fig. 4). Together, these results show that ICS imaging intrinsically encodes the information needed to distinguish localization-associated protein phenotypes, and that image-derived metrics provide a framework for translating visual variation into quantifiable features.

### Machine-learning-guided design of cell sorting strategies

FACS sorting requires predefined combinations of gating parameters and decision thresholds that define cutoffs separating target subsets from the remaining population, a process resembling a decision tree (Fig. 2f). This structure is well suited to machine learning: given a labelled target population, the optimal parameter and threshold combinations can be learned automatically. Our dataset of 50 protein tags, each profiled by ICS, provides a comprehensive representation of subcellular localization patterns for training such models. Existing automated gate-finding methods have primarily focused on isolating single pure subpopulations^41,42^, and are therefore unsuited to resolving many phenotypes at once, furthermore they have not been validated on image-derived ICS parameters. To inform sort decisions across more diverse phenotypic classes we therefore developed a progressive training framework for complex, hierarchically structured data. In three steps, it (i) computes a similarity matrix across training tags from their ICS profiles; (ii) iteratively groups tags into clusters at varying resolution, training a decision tree at each level to capture both coarse- and fine-grained distinctions; and (iii) prunes redundant trees and branches, aggregating the informative splits into an ensemble of gating rules defining optimized gating strategies (Fig. 3a).

**Figure 3:**
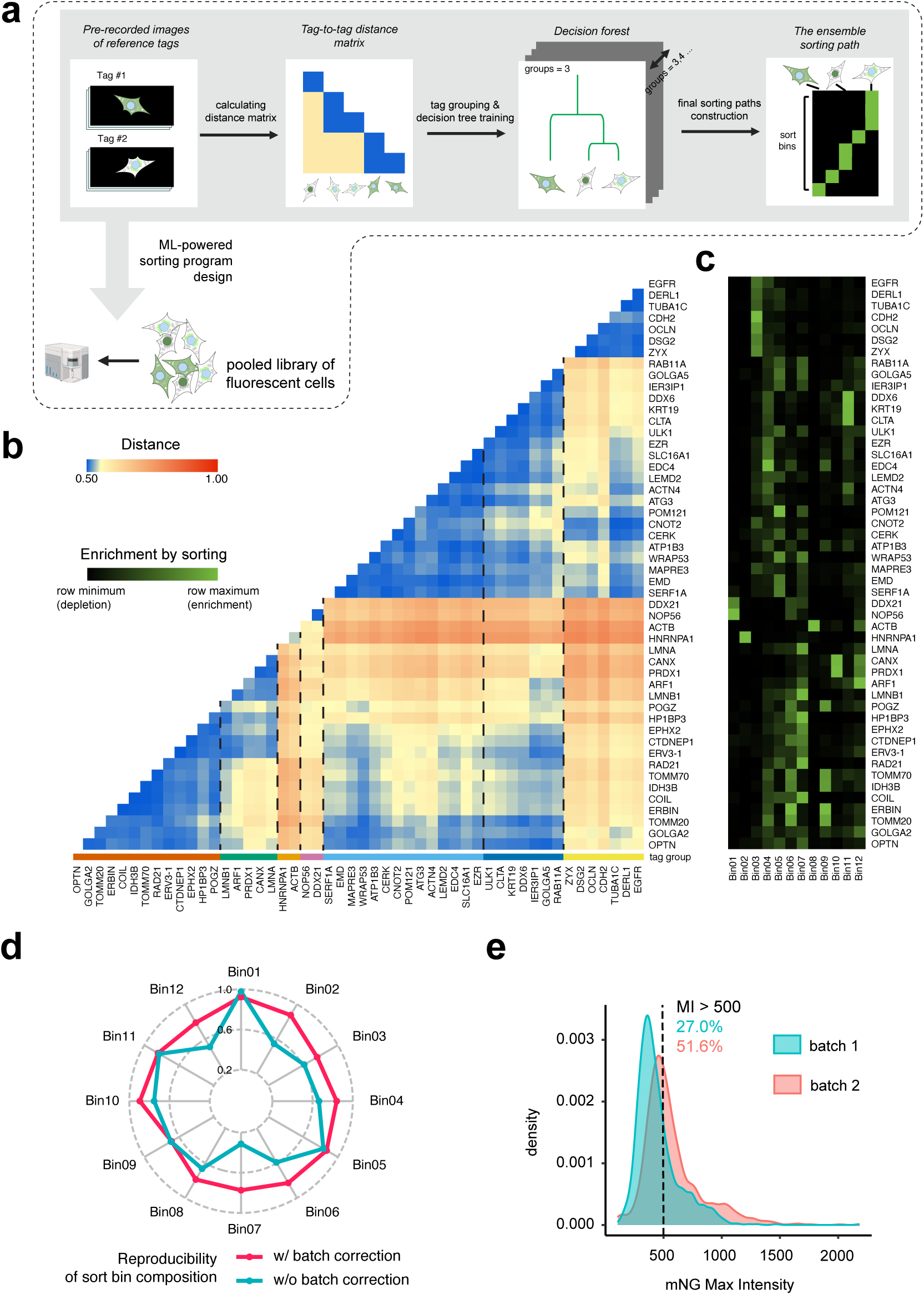
Machine-learning-guided design of sorting strategies enables the isolation of protein phenotypes. (a) Workflow for constructing sorting paths for a tagged protein library. Recorded imaging data from the training tag set was used to construct a similarity matrix, defining tag groups for decision tree training. Decision trees were de-redundified and pruned into ensemble sorting paths. (b) Distance matrix of 50 protein tags, grouped into 7 clusters based on hierarchical clustering. (c) Simulated sorting outcome of a 50-protein library into 12 sort bins. Enrichment score represents the proportion of the initial pool sorted into each bin. (d) Reproducibility comparison of sorting outcomes and reduction of batch effects with calibration, scored by the weighted Pearson correlation between experiment and prediction. (e) Illustration of batch effect: mNG max intensity measured for two batches of PRDX1-tagged cells is shown (1,000 random mNG⁺ cells each). Dashed line denotes an example sorting threshold at MI > 500, that introduces batch variation in sorting outcome (percentage of population passing the threshold displayed in their corresponding colors).

The 50 tags yielded a similarity matrix with clear hierarchical structure (Fig. 3b) that correlated with Human Protein Atlas localization annotations (https://www.proteinatlas.org/). Grouping the tags across resolution levels (i.e., varying group numbers) and training a decision tree at each produced hundreds of candidate branches, each defining a distinct gating path (Supplementary Fig. 5). These gating paths spanned coarse, compartment-level splits through to gates separating closely related localizations (Supplementary Fig. 6). After pruning redundant paths, an ensemble of gating rules defining 12 sort bins remained. In an *in silico* sorting experiment simulating compositional changes within the pooled 50-line dataset, tags showed distinct distributions across the 12 bins, indicating that the bins capture features that discriminate localization patterns (Fig. 3c). As this hierarchical gating scheme is compatible with ICS, these bins can be used to physically isolate cells and enrich specific phenotypes. The predicted distribution captured 83.9% of cells in at least one bin, with per-tag capture ranging from 99.4% (NOP56) to 55.1% (ATG3) (Supplementary Fig. 7).

Finally, since binning inherently reduces information, we asked whether the 12-bin composition alone could recapitulate the full ICS phenotype space: PCA on bin composition recovered much of the structure among the 50 tags, with similarly localized proteins clustering together (Supplementary Fig. 8a). However, these PCA-based clusters do not map one-to-one onto individual protein localizations, mainly because the gating rules underlying the 12 bin sets reflect a combination of signal localization, morphology and intensity, which cannot be fully disentangled (Supplementary Fig. 8a).

### Pooled cell sorting reproduces predicted composition via threshold calibration

To test whether the predicted compositional shifts were reproducible *in vitro*, we pooled all 50 training-tag lines in equal numbers (hereafter ‘training tag pool’) and applied the 12-bin scheme on the ICS. Sorted cells were analyzed by Anchor-Seq long-read sequencing to quantify correctly integrated HDR events across all tags, and compared with the unsorted pool (’input’) to identify enrichment. Agreement between predicted and observed distributions was scored by a weighted Pearson correlation (Fig. 3d), with down-weighting low-abundance or weakly enriched proteins that are more noise-prone.

Using the original training-derived gating rules, 7 of the 12 bins showed strong agreement (correlation >0.6), but five (bin02, bin03, bin06, bin07, and bin12) reproduced poorly. We reasoned that this discrepancy likely arises from day-to-day variation in instrument calibration / performance and sample preparation^43,44^. Indeed, replicate recordings of the same line on different days gave significantly different sorting outcomes at the same GFP-max-intensity threshold (Fig. 3e). Although calibration standards such as fluorescent beads are routine in microscopy and flow cytometry for stabilizing laser settings and detector gains^44,45^, no standard has been established for image enabled cell sorters to adjust the instrument to sample variation. To correct for this, we replaced static thresholds with dynamically adjusted ones: before each sorting experiment, three reference cell lines (TOMM20, NOP56, HNRNPA1) were recorded to compute day-specific shifts in ICS parameters. The gating thresholds were then dynamically adjusted by these offsets when the experimental samples were processed (see Materials and Methods). This daily calibration increased the prediction-to-observation correlation of all 12 bins to above 0.6 (Fig. 3d, red), with bins 06 and 12 nearly doubling. We provide this calibration tool, together with a UMAP visualization of all 50 training-tag populations, on an EMBL app server (https://apps.embl.de/icsview/).

Apart from the overall agreement between observed and predicted pool composition, individual tags varied in how closely they matched the prediction (Supplementary Fig. 8b). Lower consistency is generally correlated with lower read counts (e.g., LMNA and EGFR), indicating measurement noise. CTDNEP1 was a notable exception, with high read counts but a clear discrepancy between predicted and observed distributions, which we attribute to its low fluorescent intensity: mNG+ gating enriches relatively few correctly tagged cells and may instead capture low-abundance alternative tagging outcomes (e.g., off-target integrations), shifting the observed distribution.

### Sorting outcomes enable protein tag phenotypic inference beyond the training set

We next tested whether CODIAC generalizes beyond the training set. We added cell lines carrying tags absent from the training set to the training tag pool and sorted with the 12-bin scheme. The training tags retained their predicted distribution, while the unseen tags fell into distinct clusters, indicating phenotypic similarity to specific training-tag groups (Fig. 4a).

**Figure 4:**
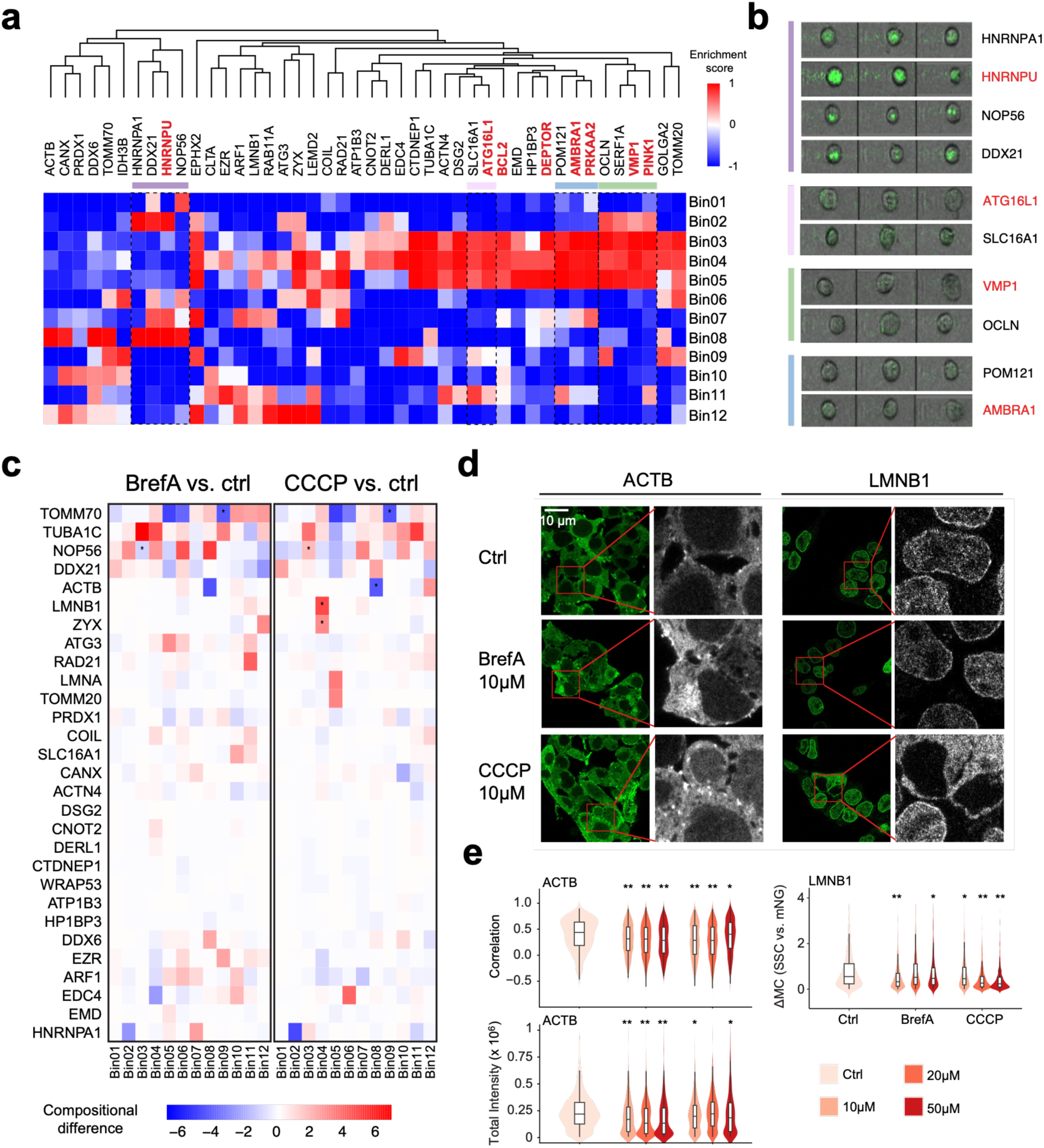
Pooled sorting with threshold calibration reproduces computational prediction and infers phenotypes of unseen tags. (a) Sorting outcome of 50 reference proteins and 7 unseen proteins. Enrichment scores are calculated based on the change in pool composition (log fold change with sigmoid transformation) relative to the initial pool. (b) Images of S8 recordings of novel cell tags compared to cells with a similar bin distribution pattern in (a). (c) Microscopy validation of the ACTB phenotype; scale bar, 20 µm. (d) Microscopy images of drug treated cells containing either endogenously tagged ACTB or LMNB1. (e) S8 recordings of ACTB-tagged cells under drug treatments at varying concentrations. For each condition and concentration, 1,000 downsampled cells are visualized as a density plot.

Proteins with similar phenotypes showed consistent bin patterns (Fig. 4a). For example; HNRNPA1 and HNRNPU, both RNA-binding members of the heterogeneous nuclear ribonucleoprotein (hnRNP) family^46^ with broad nuclear distributions, matched closely and shared enrichment in bins 2 and 8 (Fig. 4b). The nuclear proteins NOP56 and DDX21, with a granular phenotype, are positioned in the same cluster and ATG16L1 and SLC16A1 (cytoplasm and plasma membrane, respectively), with more diffuse patterns, likewise clustered together (Fig. 4b).

These results establish proof of concept that ICS sorting outcomes based on image-derived parameters can serve as encoders of protein phenotypes (i.e. phenotypic fingerprints) without isolating pure populations. Comparing the abundance of multiple tags across the range of sort bins offers a tool to systematically map the protein tag fingerprints, indicating that CODIAC is scalable to unencountered protein libraries across diverse physiological contexts.

### Utilizing CODIAC to screen for shifting subcellular phenotypes upon chemical perturbation

Finally, we asked whether CODIAC can detect perturbation-induced changes in protein localization and thus serve as a high-content screening tool. We treated the training tag pool with brefeldin A (BrefA) or CCCP. BrefA disassembles the Golgi and redistributes it into the ER^47^, and CCCP uncouples oxidative phosphorylation and dissipates the mitochondrial membrane potential^48^. Each condition was sorted in triplicate, with matched untreated controls and unsorted inputs collected the same day to control for batch and pool-variation effects.

Eight tags showed significant changes in at least one of the 12 bins, some across multiple bins (Fig. 4c). For example, CODIAC recapitulated known Brefeldin A-induced alterations in mitochondrial proteins such as TOMM70 (Fig. 4d & Supplementary Fig. 9). Notably, the cytoskeletal protein β-actin (encoded by *ACTB*), enriched in bin 8 under normal conditions, was depleted from bin 8 by BrefA or CCCP. In a concentration series on *ACTB*-tagged cells, both BrefA and CCCP shifted the mean mNG-DRAQ5 correlation and mNG total intensity (Fig. 4e), mirroring the sequencing result (Fig. 4c) and becoming more pronounced at higher dosage. Confirmed by microscopy, a subset of BrefA-treated ACTB-tagged cells accumulated fluorescent signals in perinuclear regions without nuclear overlap alongside an overall drop in signal intensity, departing from the uniform cytoplasmic distribution in untreated control cells (Fig. 4d & Supplementary Fig. 9). CCCP likewise reduced the ACTB signal and lessened its separation from the nucleus. The perinuclear BrefA phenotype is consistent with reported ACTB relocalization to the Golgi and BrefA-induced actin remodelling^49,50^, whereas CCCP promotes actin polymerization into cloud-like structures around mitochondria^51,52^.

Similarly, CCCP treatment enriched lamin B1 (encoded by *LMNB1*) in bin 4 (Fig. 4e). This may reflect depolarization-induced lamina disassembly, as lamins are caspase substrates during apoptosis^53,54^, while mitochondrial stress induces PKCδ/caspase-3–dependent lamin B1 modification and nuclear lamina disassembly^55^. Because this effect was not observed with BrefA, it is likely specific to mitochondrial depolarization rather than a general consequence of ER or secretory stress.

We reasoned that although reconstituted ICS images are of lower resolution than conventional microscopy, their far higher throughput (i.e. sampling hundreds of thousands of cells per population) combined with the CODIAC framework provides sufficient sensitivity to detect subtle shifts in fluorescence signal.

## Discussion

Measuring the spatial organization of the proteome at scale is a fundamental challenge, because protein function depends intrinsically on spatiotemporal distribution within the cell^56^. CODIAC repurposes ICS from a tool for isolating pure populations into a quantitative assay of subcellular organization. It converts spatial, morphological information into a quantifiable, scalable data type, turning spatial proteome into measurable metrics of phenotypes. This approach addresses a limitation common to existing scalable methods for spatial proteomics. Microscopy-based atlases resolve localization at high resolution, but require extensive manual curation to reach proteome scale^2,57^. Organelle profiling achieves this at the expense of single-cell resolution and with limited resolution of proteins distributed across multiple compartments^58^. Pooled image-based screens that genotype cells *in situ* rely on a terminal readout that does not recover the profiled cells^13^. Pooled multicolour-tagging approaches follow protein dynamics in live cells, but resolve identity optically, leaving throughput set by microscopy. CODIAC instead profiles many proteins in parallel from a single pooled, sequencing-based measurement, and the gating rules, once trained, are applied unchanged to new pools and conditions.

Like any instrument that images fluorescently tagged protein properties, ICS does not capture spatial distribution in isolation, but via three properties of the fluorescent signal: localization, abundance and morphology. These co-vary within a cell and cannot be disentangled from a single measurement. Whereas conventional cytometry reports abundance, size and granularity alone^44^, morphology-aware approaches such as ICS shift the emphasis toward spatial and morphological variation^26,33–36^, though abundance variation is never fully removed, given the intrinsic link between the three. Within the CODIAC framework, the balance between them is shaped by the composition of the training pool. Our proof-of-concept relied on proteins of medium-to-high expression levels with each localization represented by several proteins, a design that downweights abundance variation between proteins and makes morphology the more discriminating signal. A training set spanning a wide range of expression levels would, in principle, let the gating rules weight morphology and localization more heavily and reduce the leverage of abundance differences, at the cost of additional optimization. Nevertheless, suppressing abundance entirely would be undesirable, as it would mask drug-induced changes in protein levels^59–61^ that the framework is designed to detect.

CODIAC operates in two distinct phases: a training phase, in which a gating rule set is learned from a training tag panel, and an application phase, in which that fixed rule set is used to profile a pool of interest. Because these phases are separable, a rule set trained once can be re-used across many specific applications simply by changing the pool of interest and the test conditions. This would allow, for example, the localization changes of proteins in a given pathway to be probed by designing a dedicated pool and profiling it across perturbation conditions. Ideally, training and application are carried out in a matched cell context (e.g., the same cell type), so that the tagged protein and external perturbation remain the only two variables in the assay. Instrument-derived technical variation is unavoidable, but can be partially corrected by our calibration approach (Fig. 3e). Where the training panel must be rebuilt (e.g., when moving to a different cell background), our machine-learning framework provides a data-driven strategy for gate discovery that adapts to the new context.

A current limitation of this study is that the cell pools were assembled from individually generated CRISPR-tagged cell lines. In principle, CODIAC is compatible with pools produced by either arrayed or pooled editing, and pooled editing would offer the greater gain in scalability, since a single gating rule set can be applied to any pool once defined. Realizing this in practice, however, remains challenging.

CRISPR editing efficiency and fidelity remain limited in mammalian systems^62^, and imprecise repair outcomes generate unintended integration products that can confound insertion-site sequencing^63–65^. Improving editing efficiency and the success rate of endogenous tagging in a pooled format will therefore be needed to extend CODIAC to higher throughput.

Overall, CODIAC enables rapid, scalable, and cost-effective monitoring of protein localization and abundance across complex cellular populations while preserving the ability to physically isolate phenotypically defined cells. By bridging the throughput of flow cytometry with the spatial information of imaging, CODIAC provides a powerful new approach for functional genomics, perturbation screening, cell biology, and drug discovery applications.

## Methods

Cell culture

HEK293T cells^38^ were cultured in DMEM high glucose (Life Technologies) supplemented with 10% (vol/vol) FBS (Gibco). All cells were cultured at 37°C with 5% CO_2_ and regularly tested for mycoplasma contamination.

Cell Line Generation

### Generation of Plasmids and PCR cassettes

For tagging, the M1 and M2 tagging oligos used for generating PCR cassettes were designed using the online tool (www.pcr-tagging.com;^37^) and obtained from Sigma-Aldrich with RP1 cartridge purification. The PCR cassettes were amplified using pMaCTag-2A-N07 as template^38^ based on our previously published protocol^37^. All primers and oligos used for cloning, validation and PCR cassette generation are listed in Table S1.

For cell line generation, cells were transfected with individual PCR cassettes followed by selection as described in Lou et al., 2025^38^.

Briefly, cells at 80% confluency in one well of a 24-well plate were transfected with 100 ng Cas12a helper plasmid pVE13300 (Addgene #137715) and 900 ng of the PCR cassette template. One day after transfection, cells were expanded to a 6-well plate. Two days after transfection, cells were selected with 500 µg/ml neomycin (Sigma-Aldrich). The cells were kept in media containing neomycin for at least two weeks to remove non-edited cells from the population.

### Drug Treatments

For investigating effects of drug treatments on individual tagged proteins, cells were seeded in 8-well chamber slides and incubated overnight before drug treatment for 4 hours. Unless otherwise stated CCCP (Sigma-Aldrich, C2759) was used at 10µM and BrefA (Invitrogen, B7450) was used at 5µM. All drugs were diluted to a 1000x stock solution in DMSO (Invitrogen^TM^, D12345).

## Cell Analysis

### Fluorescence microscopy

For assessing mNG signal localization, cells were seeded in 8-well chamber slides (μ-Slide 8 Well ibiTreat 80826, ibidi) and incubated for at least 4 hours before imaging and stained with Hoechst 33342 (4 µg/ml in PBS, Thermo Fisher Scientific) for 10 min. Images were acquired on a Nikon Ti-E wide field epifluorescence microscope with a 60x ApoTIRF oil-immersion objective (1.49-NA(numerical aperture); Nikon), an LED light engine (SpectraX, Lumencor), a 2048 x 2048 pixel (6.5 µm) sCOMS (scientific complementary metal-oxide semiconductor) camera (Flash4, Hamamatsu) and an autofocus system (Perfect Focus System, Nikon) with either bright field or 469/35-nm excitation filter (Semrock) and 525/50-nm emission filter (Chroma). Z-stacks of 11 planes with 0.5-µm spacing were recorded with 100-ms exposure time for all cells, maximum intensity z-projections are shown. Subcellular localizations were identified and scored visually. For cell counting, random fields of view were inspected in the Hoechst channel, and all nuclei present in the entire field were counted. Cells were fixed using formaldehyde (ThermoScientific^TM^, 28906) at a final concentration of 4% for 5 minutes and washed twice with PBS before imaging on a Lightsheet Leica TCS SP8 DLS using a 63x oil immersion lens. Image Analysis was conducted using Fiji version 2.1.0/1.53c.

### Imaging Flow Cytometry

Cells were harvested, stored on ice prior to analysis and filtered through a 35 μm cell strainer to avoid clumping. Cells were sorted into 5 ml FACS tubes containing 80% EtOH for fixation in order to avoid degradation of genomic DNA of already sorted cells while the sort is still ongoing. Both sample and sorted tubes were kept at 4C at all times. Sorting was conducted on a BD S8 with a 100 μm sort nozzle. Viability staining with DAPI was used to sort out dying cells. DRAQ5 was added to visualize nuclei at a 1:500 concentration. Data was analyzed using FlowJo_v10.9.0. An example of gating strategy used for FACS and ICS-based sorts is added as Supplementary Fig 2.

From each batch of cells used for sorting, an input sample that underwent the same sorting procedure as bin-specific samples was collected. All samples were collected by centrifugation for 10 min at 500 × *g* at 4 °C. Pellets were frozen at −20 °C until gDNA preparation.

### Genomic DNA isolation

Genomic DNA (gDNA) was extracted from samples using Monarch Genomic DNA purification kit (NEB, T3010) according to the manufacturer’s protocol. Briefly, sorted cell pellets were resuspended in 100 µl of cold PBS and incubated with 100 µl of Cell Lysis Buffer containing Proteinase K and RNase A at 56 °C for 5 minutes. The lysates were then mixed with 400 µl Binding Buffer and loaded onto gDNA Purification Columns to bind the gDNA. The flow-through was discarded, and the columns were washed twice with 500 µl of Wash Buffer. Finally, the gDNA was eluted in 45 µl of preheated (60 °C) Elution Buffer. After elution, the gDNA concentration was measured with NanoDrop, and its integrity and size were assessed by gel electrophoresis on 1% agarose/TAE.

### Anchor-Seq library preparation and ONT sequencing

Sequencing libraries for Anchor-Seq were prepared following our previously published Anchor-Seq protocol with some modifications^16,37^. In brief, 300 ng of Tn5 (E54K, L372P, 50 ng/µl) transposase^66^ was preloaded with 1.5 µM annealed adapters (P5-UMI-xi5001…5096-ME.fw, Tn5hY-Rd2-Wat-SC3) in 50 mM Tris-HCl (pH 7.5) at 23 °C for 1h. The preloaded transposase was mixed with 400 ng of gDNA, 10 mM Tris-HCl (pH 7.5), 10mM MgCl2, and 25%[v/v] dimethylformamide, and incubated at 55 °C for 10min to tagment the gDNA. Tagmented gDNA was purified using NGSPrep beads (Steinbrenner, MDKT00010005) according to the manufacturer’s instructions. The purified tagmented gDNA was used as a template for a first PCR reaction with a biotinylated cassette-specific primer and Tn5 adapter-specific primer, using LongAmp Hot Start Taq2 polymerase (NEB, M0533). The reaction was run for 15 cycles with 63 °C annealing and 70 s elongation. Biotinylated amplicons were initially purified using NGSPrep beads, followed by enrichment with Dynabeads MyOne Streptavidin C1 beads (Invitrogen, #65001). These beads were used as the input of a second PCR, which used nested cassette-and Tn5 adapter-specific primers with LongAmp Hot Start Taq2 polymerase. The following program was used: 3 min at 95 °C; 5 cycles of 30s at 95 °C, 60 s at 60 °C, and 70 s at 65 °C; 30 cycles of 15s at 95 °C, 15 s at 62 °C, and 70 s at 65 °C; 5 min at 65 °C; and 4 °C hold. PCR products longer than 500bp were size-selected using NGSPrep beads and subsequently ligated with ONT sequencing adapter (ONT, SQK-LSK114) to generate the final library. Libraries were quantified using Qubit, loaded onto R10.4.1 flow cells according to the manual, and run on the MinION device (ONT). The primers used for Anchor-Seq are listed in Table S1.

For drug screening experiments, library preparation and sequencing were carried out as follows: Resuspended lyophilized oligonucleotides from Anchor-seq protocol^16,37^ was used for UMI tagmentation barcoding of each gDNA sample. Tn5-Rd2-Wat-SC3.rv (stock concentration 100 µM) was resuspended in Tris (stock concentration 10 mM, pH8) and distributed into 96-well plates with P5-UMI-xi50nn-ME.fw at a working concentration 50 µM for annealing. Annealed oligonucleotides were diluted to 35 µM with nuclease-free water. Tagmentation and PCR1 were performed using as previously described^66^. 2.5 µg of in-house produced Tn5E54K,L372P (25 mM Tris pH 7.5, 800 mM NaCl, 0.1 mM EDTA, 1 mM DTT, 50% glycerol) was mixed with the annealed oligonucleotides. The linker-Tn5 mixture was then incubated at 23 ℃ with constant shaking of 350 rpm for 45 min. The preloaded Tn5 was mixed with 30-100ng gDNA in 10 mM Tris-HCl pH 7.5 (Sigma Aldrich), 10 mM MgCl2 (Sigma Aldrich), and 25% dimethylformamide (DMF) (Sigma Aldrich). The reaction mixture was incubated at 55 ℃ for 10 mins. Tagmented gDNA was purified using an equal volume of SPRIselect beads (Beckman Coulter).

The purified tagmented gDNA was then used as a template for the first PCR reaction as described above. The purified PCR products were checked for size distribution using the Qubit Fluorometer and the Agilent TapeStation System and used as the input for the second PCR under the same conditions as listed above. PCR products longer than 250 bp were size-selected using SPRIselect beads and subsequently ligated with ONT sequencing adapter (ONT, SQK-LSK114) to generate the final library. The purified PCR products were then checked again for size distribution using the Qubit Fluorometer and the Agilent TapeStation System. The final libraries were loaded onto R10 nanopore flowcells (PromethION Flow Cell (FLO-PRO114M)) or MinION/GridION Flow Cell (FLO-MIN114)) and run using GridION system (ONT) following the manufacturer’s instructions.

## Computational Analysis

### Processing of ICS pre-recorded dataset

The ICS recording data was exported in .fcs and .cvw formats. The .fcs files were re-imported into FlowJo to reconstruct the workspace and apply quality filters to identify mNG⁺ DAPI⁻ cells. The resulting .wsp files, together with the corresponding .fcs files, were then processed using a custom Python script to generate a cell-by-parameter table in .tsv format for each tag population.

### Calculation of tag-to-tag distances

To compute the pairwise distances between protein tags, a dataset containing all 50 training tags was utilized. For each tag, 1,000 downsampled cells that passed the mNG-positive threshold were included in the analysis. Pairwise distance between protein tag and was calculated using the following formula:

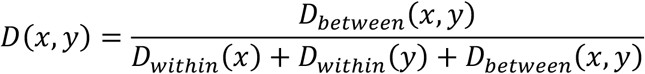

where *D*_*within*_ is defined as:

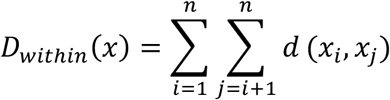

And *D_bet_*_w*een*_ is defined as:

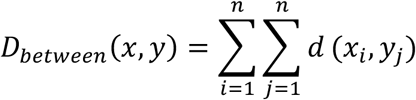

Here, *d*(*x*, *y*) represents the distance in a two-dimensional space derived from the first and second principal components of the PCA analysis using the input dataset.

### Decision tree construction and branch selection

Training tags were pre-grouped into *k* clusters prior to decision tree construction to derive distinct parameter and cutoff combinations. The value of *k* was iteratively varied from 2 to 40 using two clustering approaches: hierarchical clustering and k-means clustering. In addition, two alternative groupings based on subcellular localization of the training tags were included, resulting in a total of 80 grouping schemes.

For each grouping, tag-IDs were replaced by group IDs, which were used as labels for model training and performance evaluation. The recursive partitioning and regression tree (rpart) approach^67^ was used to train decision trees to best distinguish labelled populations within each prediction task. The data was split into training and validation sets at a 3:1 ratio, and model performance was estimated using 5-fold cross-validation repeated three times. The final model was then trained on the full training set for downstream analysis.

Each node in the decision trees represents a combination of parameters and cutoffs that mimics the ICS process, enabling prediction of compositional shifts from an initial balanced 50-tag population after sorting. The predicted post-sorting compositions were aggregated across 80 decision trees derived from 80 groupings, and a correlation matrix was computed to identify nodes with similar profiles, which were considered redundant. From this combined decision forest, an initial set of 10 nodes (i.e., sorting bins) was selected by removing nodes with pairwise correlation > 0.55. Subsequently, a core set of protein tags representing the full training space was defined by further removing redundant protein tags with correlation > 0.75. Following these steps, two additional sort bins (bin 11 and 12) were manually chosen from the remaining nodes to increase complexity of the selected sort bins.

### Gini score computation

Gini index evaluates the performance of classification models in partitioning a population based on a variable of interest. Specifically, given a classification of protein tags into classes, and a binary variable indicating a protein localization label (e.g., nuclear), the Gini score *G_L_* is defined as:

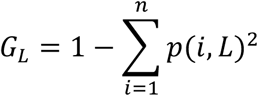

where *p*(*i*, *L*) represents the proportion of protein tag in class *i* having a localization *L*. The difference between the observed *G_L_* and a baseline *G*_O_, assuming proportional assignment of tags to classes, is expressed as:

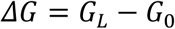

This formulation quantifies the deviation from the baseline, providing a measure of classification performance.

A permutation test was conducted for each calculated *ΔG*, with 1,000 permutations performed per test. The resulting distributions were used to determine the percentile rank of the observed *ΔG* within the corresponding random distribution.

### Assessing sort reproducibility using weighted correlation coefficients

To quantify the consistency of tag abundance between experiment *X* and prediction *Y*, a weighted Pearson’s correlation coefficient was used. For a given sorting bin, the baseline input abundance *e_i_* was measured for tag *i*. Let *x_i_* and *y_i_* denote the enrichment scores of tag *i* after sorting. The predicted enrichment is derived from the initial input abundance in the sorted sample. The weighted Pearson’s correlation coefficient between two conditions is defined as:

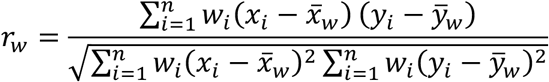

where *w_i_* is the weight assigned to the *i*-th tag, calculated as:

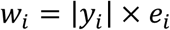

Here, *x̅_w_* and *y̅* represent the weighted means of *x_i_* and *y_i_*, respectively, and are computed as:

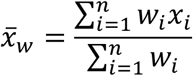

The statistic accounts for the varying importance and reliability of different tag measurements when assessing the bin of interest. Specifically, it mitigates the impact of high noise in fold changes for lowly expressed tags, ensuring a more robust measurement of consistency and reproducibility.

### Batch correction for ICS thresholds

For each ICS parameter used for sorting, baseline reference values were derived from three marker protein tags in the training dataset (TOMM20, NOP56, and HNRNPA1) by calculating the median within mNG⁺ DAPI⁻ populations. For each subsequent sorting experiment, three corresponding isolated tag samples were additionally measured on the S8 instrument on the same day, serving as calibration controls. Specifically, median values for all ICS parameters were computed for each of the three proteins and compared against the baseline reference values, with their differences capturing batch-to-batch variation between the experimental run and the training dataset.

To correct for this variation and adjust sorting thresholds accordingly, we developed a rescaling approach based on the three calibration tags. For each parameter, calibration was performed by linearly mapping values from the training reference range, defined by the minimum and maximum of the three median baseline values, to the corresponding range observed in the calibration measurements. When specified, both reference and observed values, as well as the input values, were transformed into log10 space prior to rescaling, and the calibrated values were subsequently back-transformed to the original scale. This calibration procedure was implemented in a web-based utility developed on our ShinyApp (https://apps.embl.de/icsview/) to enable automated processing.

### Integration outcome analysis

The raw signal data were generated in Pod5 formats and base-called to FASTQ files using Dorado v0.8.1 (ONT). Quality assessment of the reads was performed with NanoPlot v1.41.6. Reads were then processed using custom scripts in Julia v1.8.3 with the BioSequences v2.0.1 package for demultiplexing by barcodes and removal of Tn5 adapter and constant sequences. The demultiplexed and trimmed reads were first aligned in ONT mode using bwa-mem v0.7.17-r1188 against a custom reference genome^68^. This custom reference consisted of both wild-type and PCR cassette sequences for all target genes, with the wild-type sequence defined as up to 10,000 bases (depending on the gene length) upstream of the 5’-homology regions for each target gene. The aligned reads were filtered and annotated as HDR and insertion-deletion (INDEL) reads using in-house Perl scripts. Identified integration outcomes were evaluated using IGV v2.16.1 and further quantified by custom R scripts. The computational pipeline for characterizing and quantifying genomic integration outcomes is available on GitHub at https://github.com/dlou2022/AnchorSeq_ICS.

### Drug perturbation effect analysis

To assess the effects of perturbations on sort-bin composition, ICS sorting was performed for both treatment conditions (BrefA and CCCP) for a given bin on the same experimental day. TOMM20, NOP56, and HNRNPA1 were recorded in parallel to calibrate sorting thresholds. Sorted samples were then processed for sequencing using the protocol described above to quantify tag abundance.

For each bin-condition pair, input samples without sorting (treated and untreated) were used to estimate baseline tag abundance, and two sources of information were used to calculate the effect of perturbation: Comparisons between sorted and input samples captured enrichment or depletion due to sorting; and comparisons between treated and untreated sorted samples reflected the effect of drug treatment.

Briefly, for a given bin *u*, the fraction of tag *X* under condition *A* was calculated, transformed into a Z-score, and capped at 10, denoted as *fr*(*u*, *X*, *A*). Input samples from the same experimental day were aggregated, and enrichment was defined as the difference between sorted and input fractions:

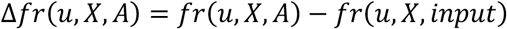

where positive values indicate enrichment and negative values indicate depletion of tag *X* in bin *u*.

To ensure robustness, a MAD-based approach was applied to filter out outlier measurements of Δ*fr*(*u*, *X*, *A*) across replicates. Measurements deviating more than five times the MAD from the median were considered outliers and excluded. The remaining data were then used to fit a logistic regression model:

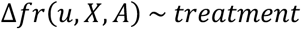

where treatment is a binary variable (treated vs. untreated).

Significant hits were defined as those with regression coefficients >1 or <−1 and a p-value <0.1, and were retained for downstream analysis.

## Code availability

ICS pre-recorded data processing and visualization: https://github.com/shli-embl/ICS_image_analysis. The long-read sequencing data analysis pipeline: https://github.com/dlou2022/AnchorSeq_ICS.

## Author Contributions

MESK, SL, MK and LMS conceived the study. MESK, SL and DL designed the study. MESK, DL, HJ, DK and KG performed experiments. DL and SL analyzed sequencing data. SL performed machine learning and computational analysis. MESK, DL and SL contributed to data visualization. All authors provided guidance and feedback on analysis and results. MESK, SL and DL wrote the manuscript with input from DS, MK and LMS and feedback from all authors. MESK compiled and edited the final manuscript. MK and LMS acquired funding.

## Acknowledgements

We thank all members of the EMBL Flow Cytometry Core Facility; Beata Ramasz, Daniel Gimenes and Alex Cabrera for help in facilitating S8 experiments and critical input. We also thank EMBL GeneCore for their help with sequencing the perturbation screen, particularly Hilal Ozgur and Vladimir Benes. We thank Fabian Mulder for help with computational data curation during early stages of the project and Rocio Nieto Arellano for help with the initial recording of the training tag library. Kai Fenzl and Maximilian Seidel provided insightfull feedback during the writing process of the manuscript. Furthermore we thank all members of the Steinmetz lab for critical input during all stages of this project. We are grateful to Simon Anders for support with the design and analysis of the sequencing experiments.

Figures 2f, 4a and Supplementary Fig. 2a were created using BioRender. License: Steinmetz, L. (2026) https://BioRender.com/r3170tv. This research was financed through a research grant by the Baden-Württemberg Stiftung (to MK and LMS) through the ‘Methoden in den Lebenswissenschaften’ Programme (MET-ID53).

## Supplementary Figures

**Supplementary Fig. 1:**
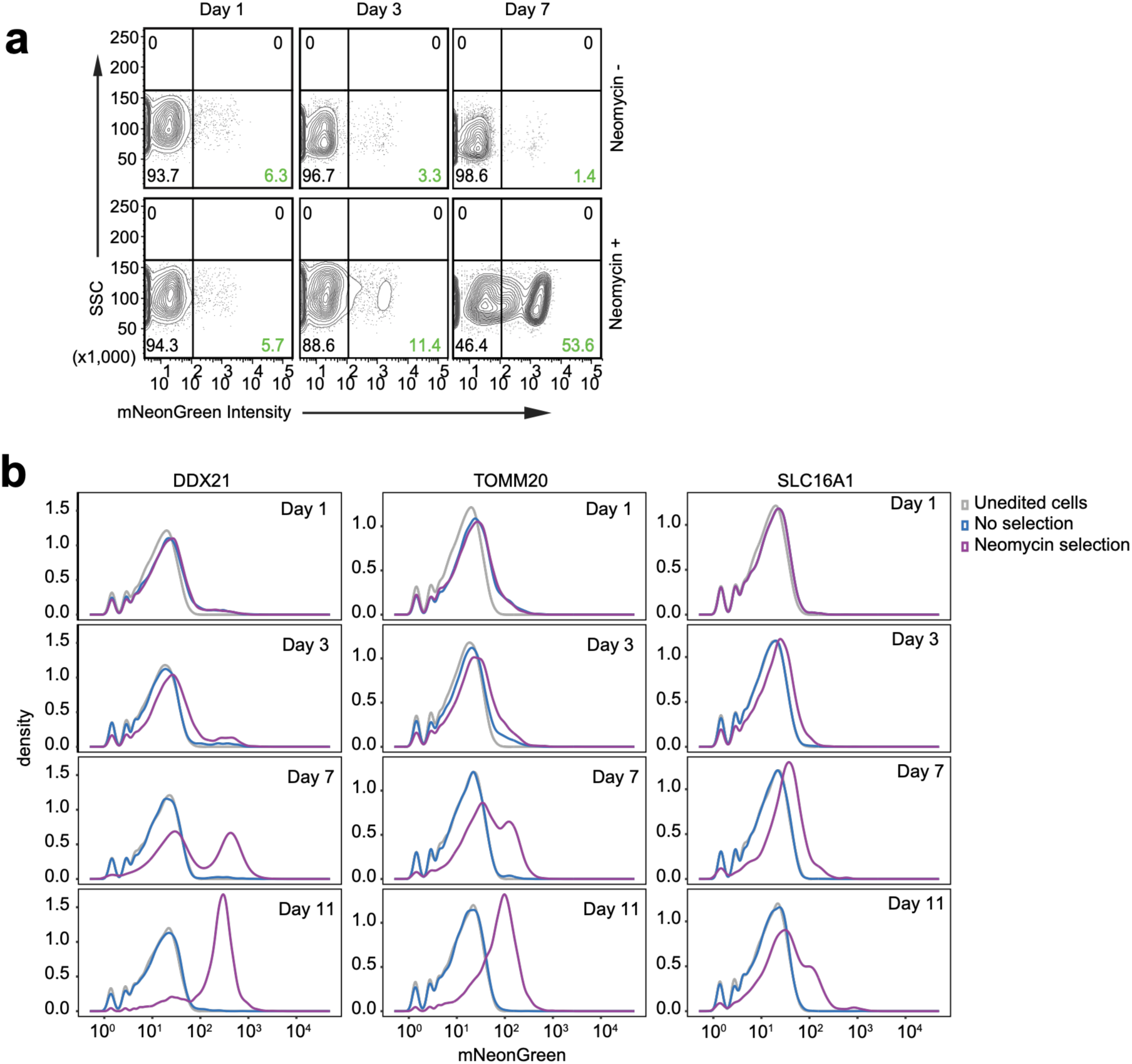
CRISPR-Cas12a tagging with Neomycin selection to enrich mNG-positive cells. a) Representative flow cytometry plots showing mNG integration into the HNRNPA1 locus in HEK293T-N cells over 7 days. b) Time course assessment of PCR tagging efficiency at multiple gene loci (*DDX21*, *TOMM20*, *SLC16A1*) in HEK293T-N cells over 11 days by flow cytometry, corresponding to Fig2b in presence and absence of Neomycin. c) Schematic representation of sequencing reads obtained by Oxford Nanopore sequencing (ONT) and integration outcome following homology-integration repair (HDR). Barcodes (BCs) were introduced during the Tn5 tagmentation step, enabling pooling of multiple samples. The HDR-mediated integration results in scarless insertion of mNG at the C-terminal of the target gene.

**Supplementary Fig 2:**
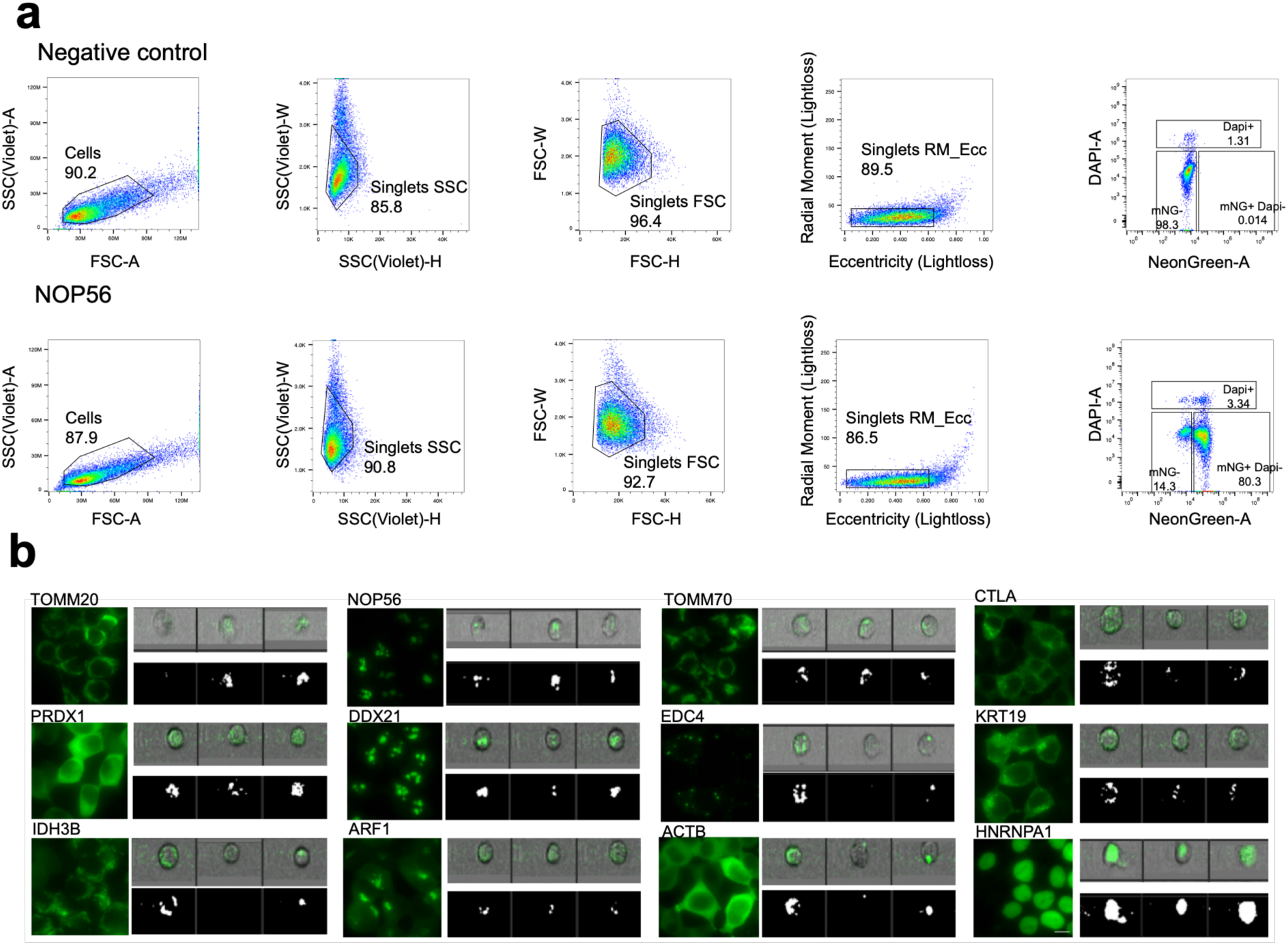
Image-enabled cell sorting gating strategy. (a) FCS plots for recording of single-cell populations on the BD S8. (b) Example images of cells tagged with mNG in microscopy (left) and S8 (right). Scale bar indicates 10μm.

**Supplementary Fig 3:**
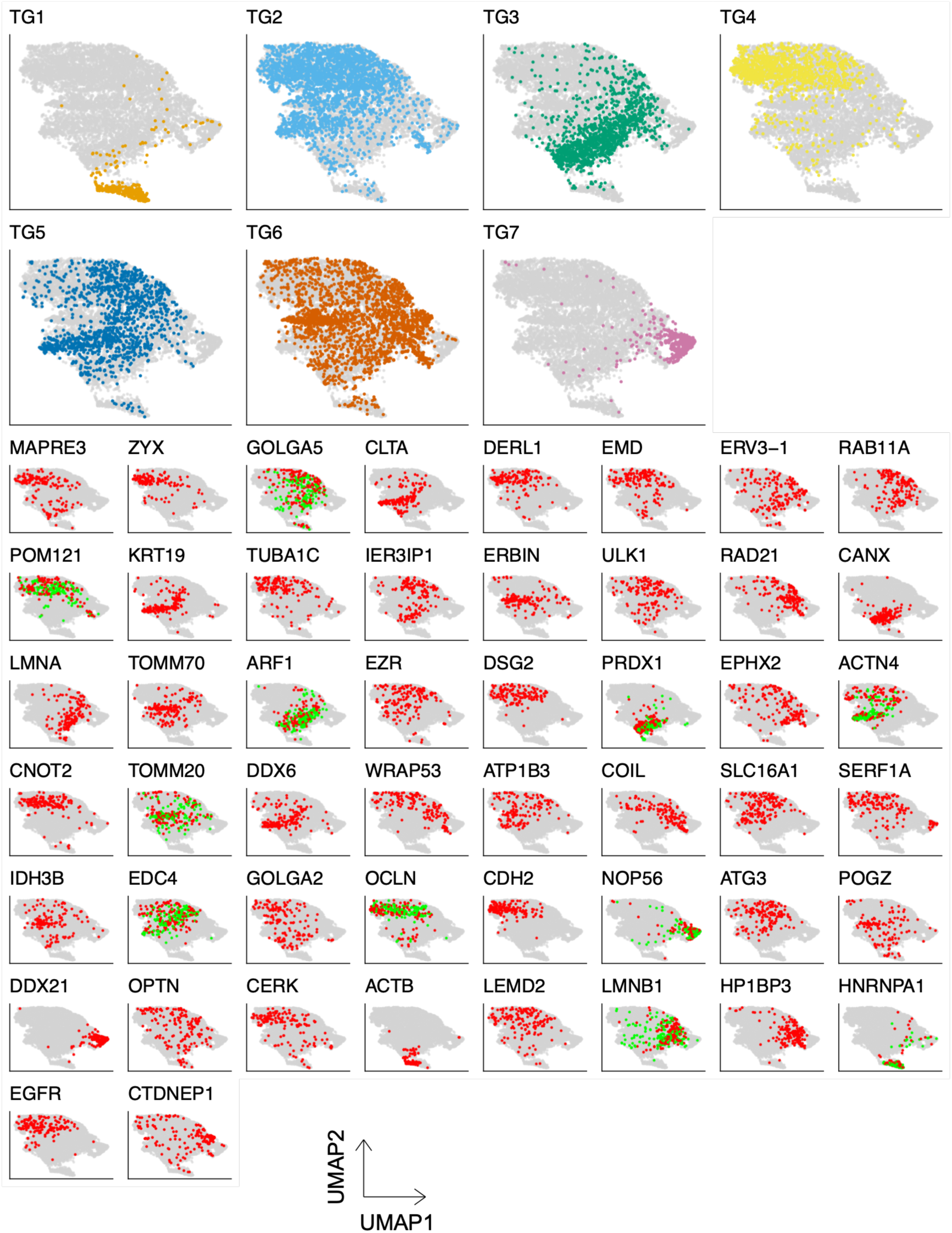
Distinct protein clusters identified via mNG-relevant features within the 50-tag dataset. Projection of tag groups (upper panel) and individual protein tags (lower panel) onto UMAP space constructed from S8 recording data using 50 training tags. For protein tags with replicated recordings, batches are indicated by colors (red: training batch; green: replicate batch).

**Supplementary Fig. 4:**
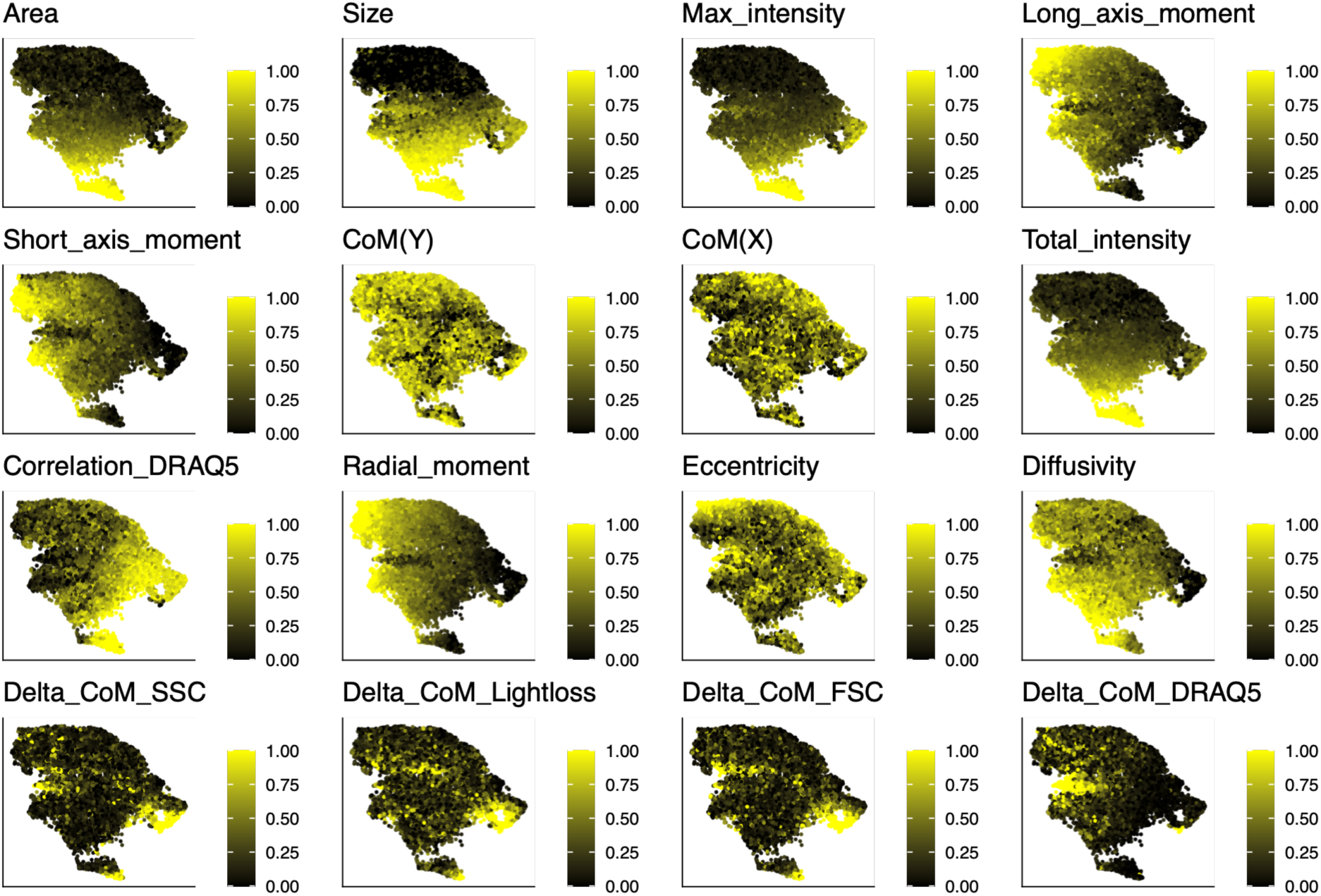
Image-derived sorting parameter profiles among 50 training tag populations. mNG-related parameter distribution across cells recorded for 50 training tags. original values were rescaled to 0-1 for visualization. CoM: center of mass.

**Supplementary Fig. 5:**
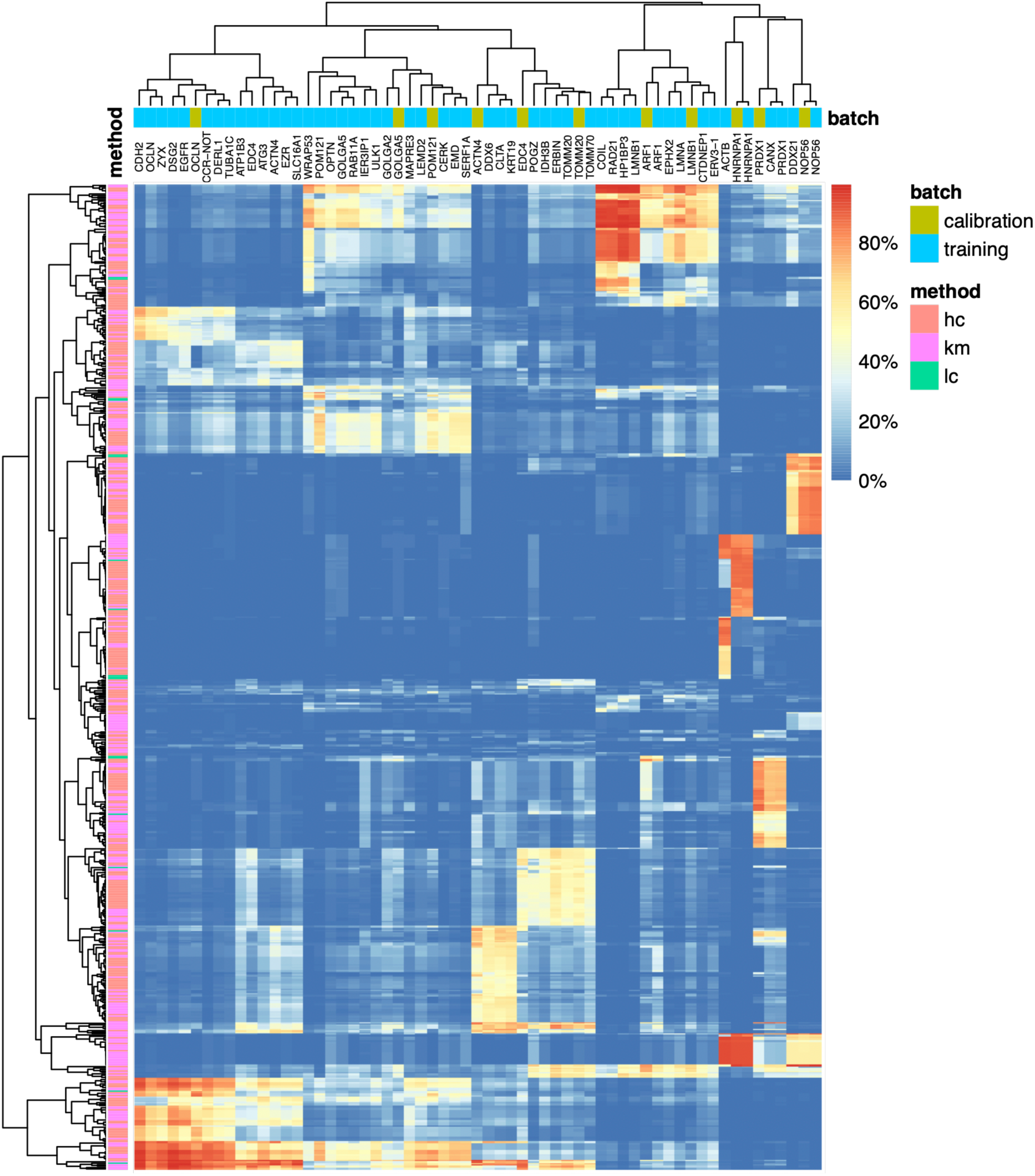
Decision forest constructed from three label-grouping approaches. Heatmap visualization of decision trees derived from 80 groupings generated using 3 methods (hc: hierachical clustering; km: k-means clustering; lc: localization-based), based on 50 training tags for RPART model training. Each row indicates a branch from a decision tree with protein tag composition predicted by applying corresponding cutoffs to the training data. Replicates of a subset of training tags (calibration batch) were included as an external validation.

**Supplementary Fig. 6:**
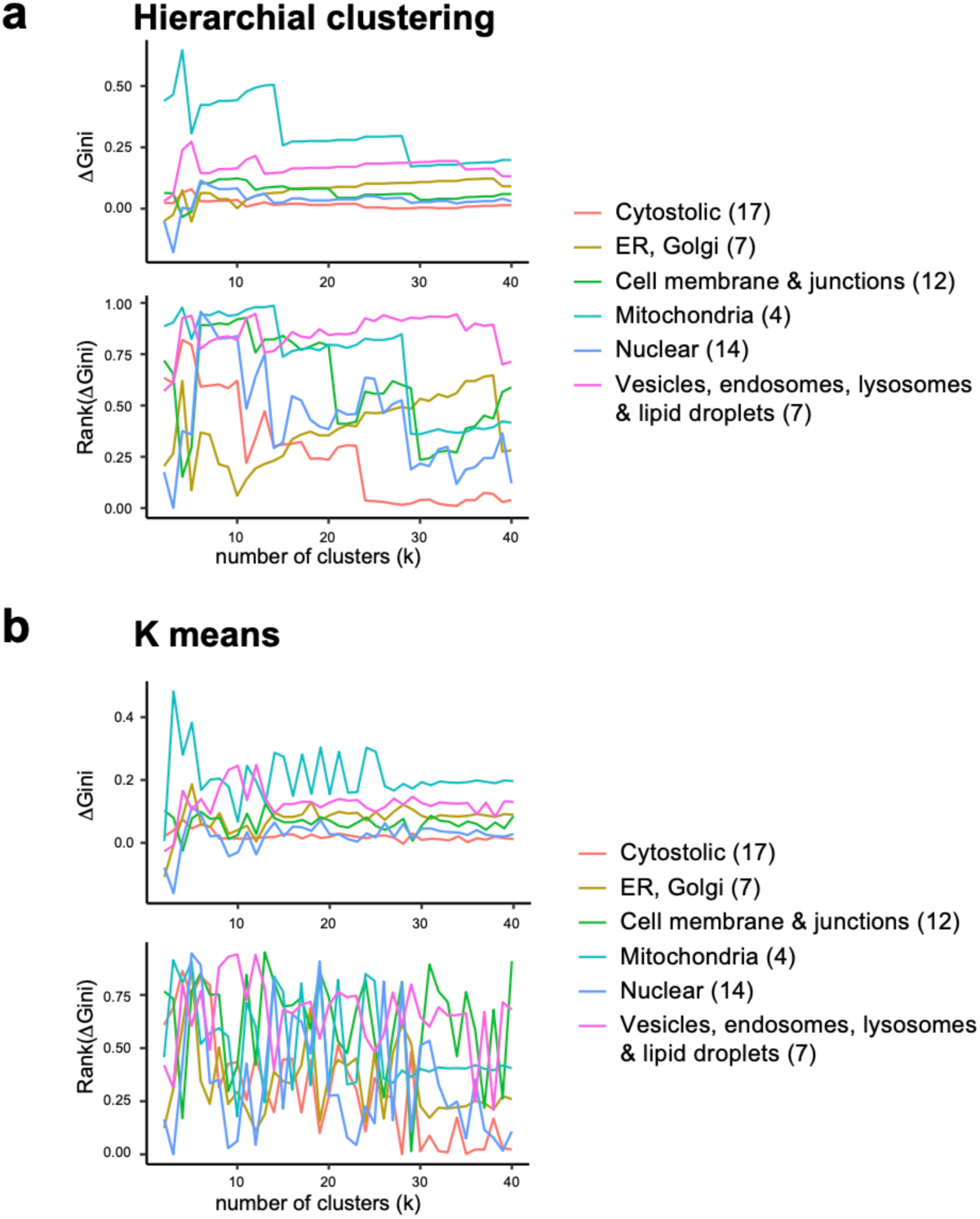
Label groupings capture protein localization distinctions assessed by Gini index. Distribution Gini index score across subcellular compartments under different cluster numbers for (a) hierarchical clustering and (b) k-means clustering. ΔGini represents the difference between the observed and a background assignment in which tags are evenly distributed into the same number of groups across localizations. Rank(ΔGini) indicates the observed classification among 1,000 random permutations with the same cluster number k, reflecting the ability to distinguish localization-specific patterns.

**Supplementary Fig. 7:**
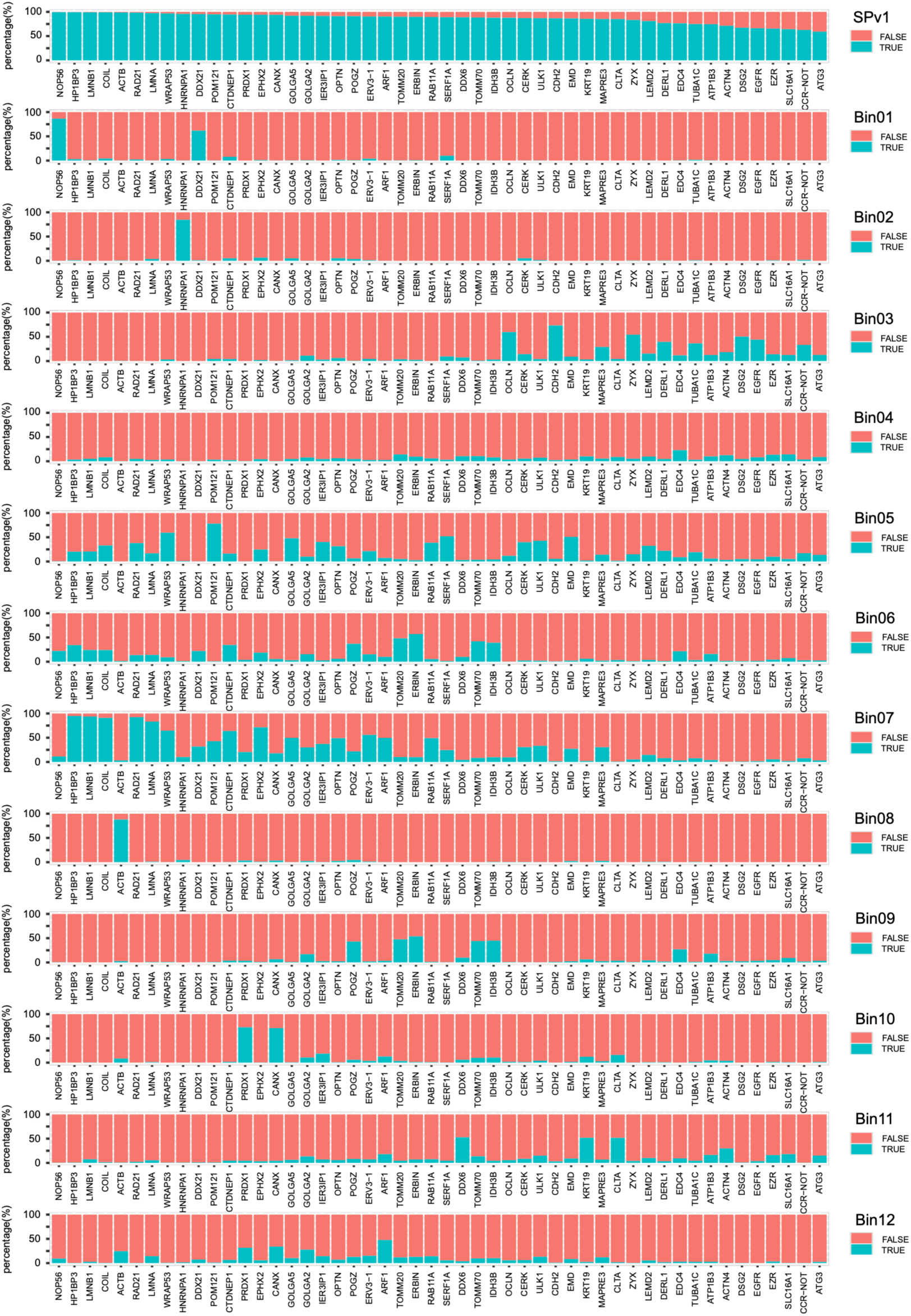
Predicted sorting outcomes of 12 bins derived from machine learning. An in silico prediction of ICS outcome for 50 training tags. Each tag was initialized with an equal number of cells. For each sorting bin, the proportion of cells qualified in each tag is shown as a bar plot. SPv1 represents the union of all 12 sorting bins, with the fraction of cells captured by at least 1 out of 12 bins indicated.

**Supplementary Fig. S8:**
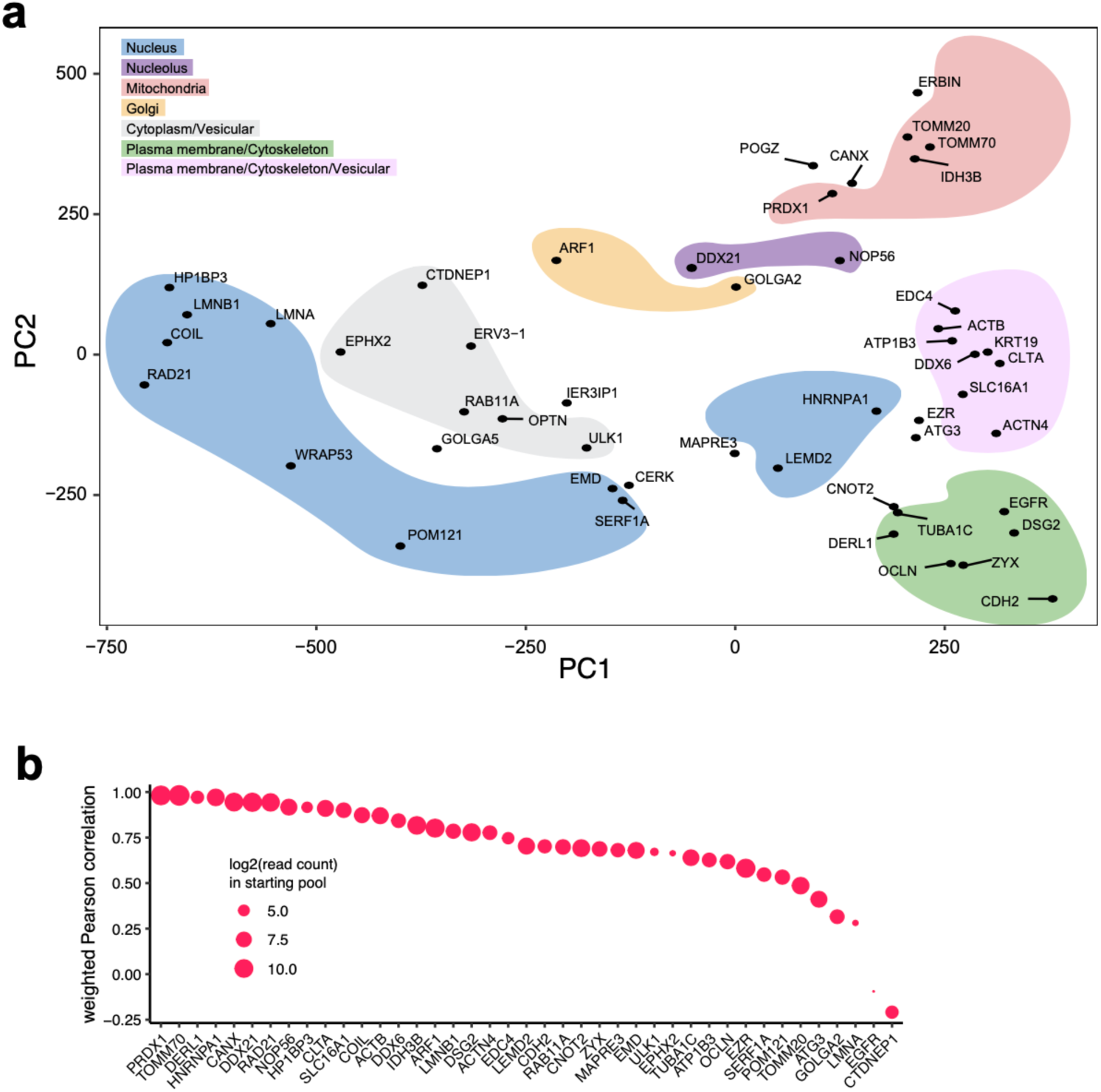
The 12-bin scheme recapitulates phenotypic diversity across 50 training tags. (a) Principle Component Analysis (PCA) computed based on ICS images and parameters of the fluorescently tagged cell lines which were simulated for their sorting outcome across the 12 sort bins. (b) The weighted Pearson correlation coefficient for 33 protein tags indicating the overlap of the biological sorting outcome with the computational prediction. The 17 protein tags not shown here were excluded due to a low read count in the initial pool.

**Supplementary Figure 9:**
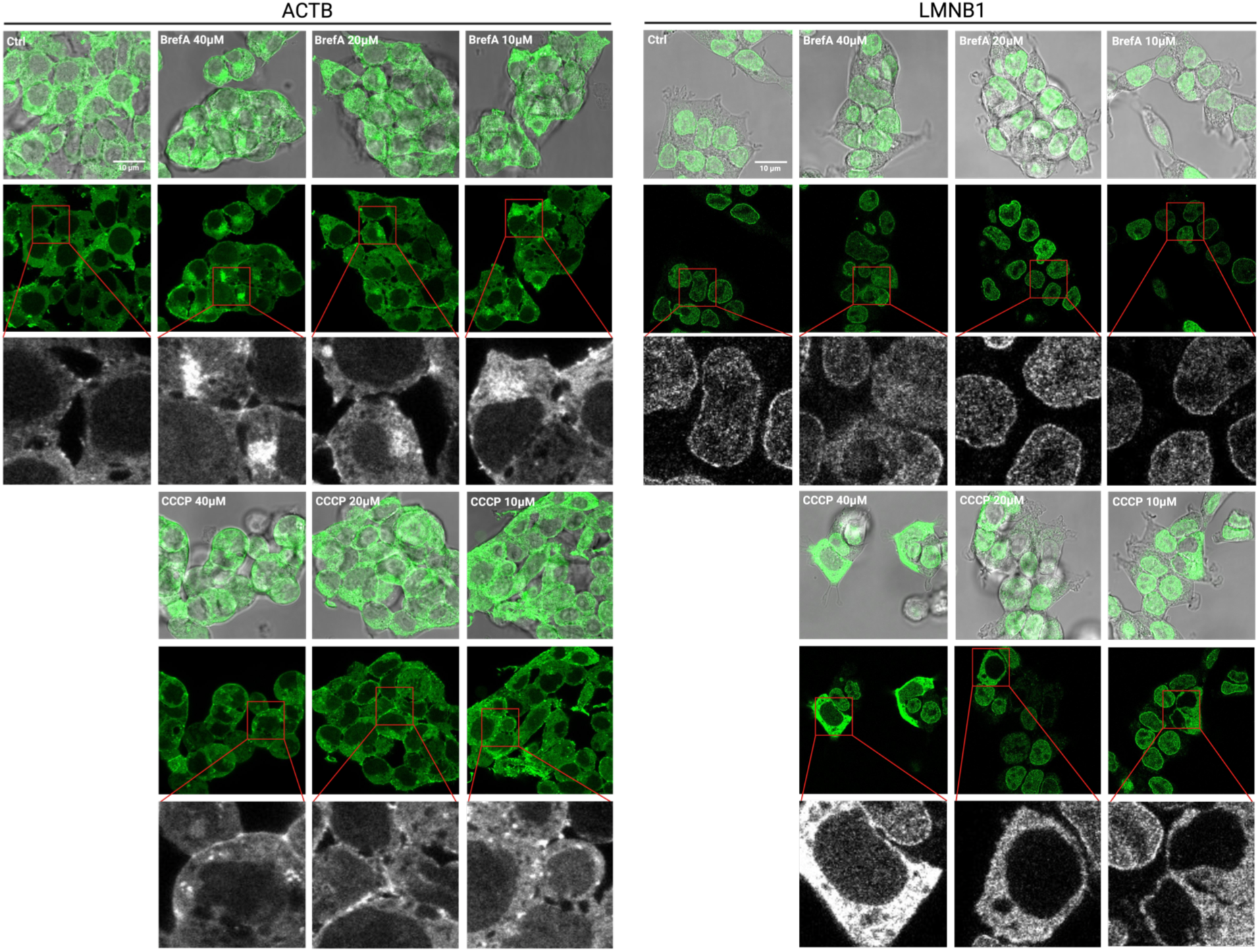
Microscopy of mNG-ACTB and mNG-LMNB1 cell lines in presence of varying screening drug concentrations. Hek293T-N cells were treated with BrefA or CCCP at varying concentrations for 4 hours, fixed with 4% FA and imaged using confocal microscopy.

## Notes

### Competing Interest Statement

The authors have declared no competing interest.

## References

1. Lacoste, J. et al. Pervasive mislocalization of pathogenic coding variants underlying human disorders. Cell 187, 6725–6741.e13 (2024).

2. Cho, N. H. et al. OpenCell: Endogenous tagging for the cartography of human cellular organization. Science 375, eabi6983 (2022).

3. Reicher, A. et al. Pooled multicolour tagging for visualizing subcellular protein dynamics. Nat. Cell Biol. 26, 745–756 (2024).

4. Nemčko, F. et al. Proteome-scale tagging and functional screening in mammalian cells by ORFtag. Nat. Methods 21, 1668–1673 (2024).

5. Leonetti, M. D., Sekine, S., Kamiyama, D., Weissman, J. S. & Huang, B. A scalable strategy for high-throughput GFP tagging of endogenous human proteins. Proc. Natl. Acad. Sci. 113, E3501–E3508 (2016).

6. Valm, A. M. et al. Applying systems-level spectral imaging and analysis to reveal the organelle interactome. Nature 546, 162–167 (2017).

7. Lin, J., Fallahi-Sichani, M., Chen, J. & Sorger, P. K. Cyclic Immunofluorescence (CycIF), A Highly Multiplexed Method for Single-cell Imaging. Curr. Protoc. Chem. Biol. 8, 251–264 (2016).

8. Breckels, L. M. et al. Advances in spatial proteomics: Mapping proteome architecture from protein complexes to subcellular localizations. Cell Chem. Biol. 31, 1665–1687 (2024).

9. Christopher, J. A. et al. Subcellular proteomics. Nat. Rev. Methods Primer 1, 32 (2021).

10. Schaffer, L. V. et al. Multimodal cell maps as a foundation for structural and functional genomics. Nature 642, 222–231 (2025).

11. Lin, C. et al. Submicrometre geometrically encoded fluorescent barcodes self-assembled from DNA. Nat. Chem. 4, 832–839 (2012).

12. Ramezani, M. et al. A genome-wide atlas of human cell morphology. Nat. Methods 22, 621–633 (2025).

13. Feldman, D. et al. Optical Pooled Screens in Human Cells. Cell 179, 787–799.e17 (2019).

14. Sansbury, S. E. et al. Pooled tagging and hydrophobic targeting of endogenous proteins for unbiased mapping of unfolded protein responses. Mol. Cell 85, 1868–1886.e12 (2025).

15. Kaufman, T. et al. Visual barcodes for clonal-multiplexing of live microscopy-based assays. Nat. Commun. 13, 2725 (2022).

16. Meurer, M. et al. Genome-wide C-SWAT library for high-throughput yeast genome tagging. Nat. Methods 15, 598–600 (2018).

17. Buchmuller, B. C. et al. Pooled clone collections by multiplexed CRISPR-Cas12a-assisted gene tagging in yeast. Nat. Commun. 10, 2960 (2019).

18. Rane, A. S., Rutkauskaite, J., deMello, A. & Stavrakis, S. High-Throughput Multi-parametric Imaging Flow Cytometry. Chem 3, 588–602 (2017).

19. Goda, K. et al. High-throughput single-microparticle imaging flow analyzer. Proc. Natl. Acad. Sci. 109, 11630–11635 (2012).

20. Miura, T. et al. On-chip light-sheet fluorescence imaging flow cytometry at a high flow speed of 1 m/s. Biomed. Opt. Express 9, 3424 (2018).

21. Holzner, G. et al. High-throughput multiparametric imaging flow cytometry: toward diffraction-limited sub-cellular detection and monitoring of sub-cellular processes. Cell Rep. 34, 108824 (2021).

22. George, T. C. et al. Distinguishing modes of cell death using the ImageStream® multispectral imaging flow cytometer. Cytometry A 59A, 237–245 (2004).

23. Nitta, N. et al. Intelligent Image-Activated Cell Sorting. Cell 175, 266–276.e13 (2018).

24. Mikami, H. et al. Virtual-freezing fluorescence imaging flow cytometry. Nat. Commun. 11, 1162 (2020).

25. Isozaki, A. et al. Intelligent image-activated cell sorting 2.0. Lab. Chip 20, 2263–2273 (2020).

26. Schraivogel, D. et al. High-speed fluorescence image–enabled cell sorting. Science 375, 315–320 (2022).

27. Wang, J., Fei, B., Geahlen, R. L. & Lu, C. Quantitative analysis of protein translocations by microfluidic total internal reflection fluorescence flow cytometry. Lab. Chip 10, 2673 (2010).

28. Muñoz, H. E. et al. Single-Cell Analysis of Morphological and Metabolic Heterogeneity in *Euglena gracilis* by Fluorescence-Imaging Flow Cytometry. Anal. Chem. 90, 11280–11289 (2018).

29. Ota S. et al. Ghost cytometry. Science 360, 1246–1251 (2018).

30. Fasler-Kan, E., Baiken, Y., Vorobjev, I.A., & Barteneva, N.S. Analysis of Nucleocytoplasmic Protein Shuttling by Imaging Flow Cytometry. in Imaging Flow Cytometry. Methods in Molecular Biology vol. 1389 (Humana Press).

31. Salek, M. et al. COSMOS: a platform for real-time morphology-based, label-free cell sorting using deep learning. Commun. Biol. 6, 971 (2023).

32. Paul, R. et al. Versatile Image-Assisted Cell Sorting by Selective Trapping with Spatiotemporal Multiparameter Targeting. ACS Sens. 10, 6714–6723 (2025).

33. Kornienko, J. et al. Mislocalization of pathogenic RBM20 variants in dilated cardiomyopathy is caused by loss-of-interaction with Transportin-3. Nat. Commun. 14, 4312 (2023).

34. Pretto, S. et al. A functional single-cell metabolic survey identifies Elovl1 as a target to enhance CD8+ T cell fitness in solid tumours. Nat. Metab. 7, 508–530 (2025).

35. Sun, J. et al. Metabolic regulator LKB1 controls adipose tissue ILC2 PD-1 expression and mitochondrial homeostasis to prevent insulin resistance. Immunity 57, 1289–1305.e9 (2024).

36. Pae, J. et al. Transient silencing of hypermutation preserves B cell affinity during clonal bursting. Nature 641, 486–494 (2025).

37. Fueller, J. et al. CRISPR-Cas12a–assisted PCR tagging of mammalian genes. J. Cell Biol. 219, e201910210 (2020).

38. Lou, D. et al. Improved cloning-free one-step CRISPR-Cas12a-assisted tagging of mammalian genes using PCR generated reagents (PCR tagging). 10.64898/2025.12.02.690677(2025) doi:10.64898/2025.12.02.690677.

39. Shaner, N. C. et al. A bright monomeric green fluorescent protein derived from Branchiostoma lanceolatum. Nat. Methods 10, 407–409 (2013).

40. Hockemeyer, D. et al. Efficient targeting of expressed and silent genes in human ESCs and iPSCs using zinc-finger nucleases. Nat. Biotechnol. 27, 851–857 (2009).

41. Becht, E. et al. Reverse-engineering flow-cytometry gating strategies for phenotypic labelling and high-performance cell sorting. Bioinformatics 35, 301–308 (2019).

42. Triana, S. et al. Single-cell proteo-genomic reference maps of the hematopoietic system enable the purification and massive profiling of precisely defined cell states. Nat. Immunol. 22, 1577–1589 (2021).

43. Finak, G. et al. Standardizing Flow Cytometry Immunophenotyping Analysis from the Human ImmunoPhenotyping Consortium. Sci. Rep. 6, 20686 (2016).

44. Cossarizza, A. et al. Guidelines for the use of flow cytometry and cell sorting in immunological studies (third edition). Eur. J. Immunol. 51, 2708–3145 (2021).

45. Wu, Y. et al. The development of quantum dot calibration beads and quantitative multicolor bioassays in flow cytometry and microscopy. Anal. Biochem. 364, 180–192 (2007).

46. Geuens, T., Bouhy, D. & Timmerman, V. The hnRNP family: insights into their role in health and disease. Hum. Genet. 135, 851–867 (2016).

47. Chardin, P. & McCormick, F. Brefeldin A: The Advantage of Being Uncompetitive. Cell 97, 153–155 (1999).

48. Heytler P.G. Uncoupling of oxidative phosphorylation by carbonyl cyanide phenylhydrazones. I. Some characteristics of m-Cl-CCP action on mitochondria and chloroplasts. Biochemistry 2:357–61 (1963) doi:10.1021/bi00902a031.

49. Alvarez, C. & Sztul, E. S. Brefeldin A (BFA) disrupts the organization of the microtubule and the actin cytoskeletons. Eur. J. Cell Biol. 78, 1–14 (1999).

50. Shen, X. et al. Brefeldin A-inhibited ADP-ribosylation factor activator BIG2 regulates cell migration via integrin β1 cycling and actin remodeling. Proc. Natl. Acad. Sci. 109, 14464–14469 (2012).

51. Moore, A. S., Wong, Y. C., Simpson, C. L. & Holzbaur, E. L. F. Dynamic actin cycling through mitochondrial subpopulations locally regulates the fission–fusion balance within mitochondrial networks. Nat. Commun. 7, 12886 (2016).

52. Fung, T. S., Ji, W.-K., Higgs, H. N. & Chakrabarti, R. Two distinct actin filament populations have effects on mitochondria, with differences in stimuli and assembly factors. J. Cell Sci. 132, jcs234435 (2019).

53. Orth, K., Chinnaiyan, A. M., Garg, M., Froelich, C. J. & Dixit, V. M. The CED-3/ICE-like Protease Mch2 Is Activated during Apoptosis and Cleaves the Death Substrate Lamin A. J. Biol. Chem. 271, 16443–16446 (1996).

54. Rao, L., Perez, D. & White, E. Lamin proteolysis facilitates nuclear events during apoptosis. J. Cell Biol. 135, 1441–1455 (1996).

55. Charli, A. et al. Mitochondrial Stress Disassembles Nuclear Architecture through Proteolytic Activation of PKCδ and Lamin B1 Phosphorylation in Neuronal Cells: Implications for Pathogenesis of Age-related Neurodegenerative Diseases. Front Cell Neurosci 10.1101/2024.11.01.621517(2025) doi:10.1101/2024.11.01.621517.

56. Thul, P. J. et al. A subcellular map of the human proteome. Science 356, eaal3321 (2017).

57. Thul, P. J. & Lindskog, C. The human protein atlas: A spatial map of the human proteome. Protein Sci. 27, 233–244 (2018).

58. Hein, M. Y. et al. Global organelle profiling reveals subcellular localization and remodeling at proteome scale. Cell 188, 1137–1155.e20 (2025).

59. Winter, G. E. et al. Phthalimide conjugation as a strategy for in vivo target protein degradation. Science 348, 1376–1381 (2015).

60. Matyskiela, M. E. et al. A novel cereblon modulator recruits GSPT1 to the CRL4CRBN ubiquitin ligase. Nature 535, 252–257 (2016).

61. Krönke J et al. Lenalidomide Causes Selective Degradation of IKZF1 and IKZF3 in Multiple Myeloma Cells. Science 10.1126/science.1244851(2013) doi:10.1126/science.1244851.

62. Villiger, L. et al. CRISPR technologies for genome, epigenome and transcriptome editing. Nat. Rev. Mol. Cell Biol. 25, 464–487 (2024).

63. Banan, M. Recent advances in CRISPR/Cas9-mediated knock-ins in mammalian cells. J. Biotechnol. 308, 1–9 (2020).

64. Khan, F.A. Biotechnology Fundamentals. (CRC Press, 2020). 10.1201/9781003024750.

65. Ali, A., Xiao, W., Babar, M. E. & Bi, Y. Double-Stranded Break Repair in Mammalian Cells and Precise Genome Editing. Genes 13, 737 (2022).

66. Hennig, B. P. et al. Large-Scale Low-Cost NGS Library Preparation Using a Robust Tn5 Purification and Tagmentation Protocol. G3 GenesGenomesGenetics 8, 79–89 (2018).

67. Therneau T & Atkinson B. rpart: Recursive Partitioning and Regression Trees. (2023).

68. Li, H. & Durbin, R. Fast and accurate short read alignment with Burrows–Wheeler transform. Bioinformatics 25, 1754–1760 (2009).

